# Explainable AI-Driven Diagnosis Model for Early Glaucoma Detection Using Grey-Wolf Optimized Extreme Learning Machine Approach

**DOI:** 10.1101/2025.05.15.654213

**Authors:** Debendra Muduli, Santosh Kumar Sharma, Sujata Dash, Bernardo Lemos, Saurav Mallik

**Affiliations:** Department of Computer Science and Engineering, C.V. Raman Global University, Bidya nagar, Odisha 752054, India; Department of Information Technology, Nagaland University, India; Department of Environmental Health, Harvard T H Chan School of Public Health, Boston, MA 02115, USA; Department of Pharmacology & Toxicology, University of Arizona, AZ 85721, USA

**Keywords:** Intraocular Pressure, Extreme Learning Machine, Improved Grey Wolf Optimization, principal component analysis, Linear Discriminant Analysis, Explainable Artificial Intelligence

## Abstract

Glaucoma is a prominent threat to vision and ranks as the third leading cause of blindness in India. Early detection is crucial to limit its progression. Retinal image analysis, particularly computer-aided diagnosis (CAD), has gained significant attention due to its potential in effectively screening for and managing glaucoma. Nowadays, Artificial intelligence (AI) has achieved significant progress has been made in medical image analysis across various applications. However, the main challenge hindering the widespread adoption of deep neural models in clinical settings is their lack of interpretability. In response to this need, we propose an enhanced CAD model with four key components: image pre-processing, feature extraction, feature dimensionality reduction, and classification. Utilizing the G1020 and ORIGA datasets, we employ a fast discrete curvelet transform with wrapping (FDCT-WRP) for curve-like feature extraction. A combined feature reduction technique, principal component analysis (PCA) and linear discriminant analysis (LDA), is applied to generate relevant features and reduce the feature dimensionality. Incorporating an improved grey wolf Optimization (IMGWO) with an Extreme Learning Machine (ELM) for classification. Then, IMGWO optimizes parameters, enhancing the efficiency of single-hidden-layer feedforward neural networks. Through a 10 × 5-fold stratified cross-validation (SCV) test on standard datasets, our CAD model demonstrates superior performance. The proposed CAD model achieved better classification result i.e., 93.87% and 95.38% on the G1020 and ORIGA datasets respectively. Additionally, we have proposed a framework named as GlaucoXAI (Glaucoma explainable artificial intelligence), which utilizes seven advanced explanation methods to improve the interpretability of deep learning models for medical experts’ trust. GlaucoXAI has been employed on glaucoma detection using fundus images. GlaucoXAI’s adaptable design could aid ophthalmologists and medical professionals in detection of glaucoma. The experimental results demonstrate that the proposed model surpasses other existing models in classification accuracy, while significantly reducing the number of features.

## 1. Introduction

Recently, in the healthcare domain, there has been an ethical issue related to transparency in AI. The lack of trust stems from the opaque functioning of AI models, which need to be simplified [1]. Explainable AI (XAI) techniques refer to the methods used to elucidate AI models and their predictions [2]. Health information gathered from individual users can be useful for ophthalmologists in identifying diseases more effectively [3]. These models enable continuous health monitoring through healthcare tools like smartphones or wearable devices, empowering individuals to manage their own health effectively. Computer-aided diagnosis (CAD) for ocular diseases because of its extensive advantages, such as enabling quick and precise large-scale screening and alleviating the workload of physicians in routine clinical task [4]. Glaucoma is a persistent eye condition marked by optic nerve (ON) damage and visual impairment, manifested through the cupping of the optic disc (OD), the degeneration of ON fibres [5, 6]. Reducing intraocular pressure (IOP) has been established as a reliable and evidence-supported approach to treating open-angle glaucoma (OAG) [7, 8]. Timely diagnosis and effective management of IOP are crucial for sustaining a high quality of life, a matter of increasing significance in contemporary ageing populations. Typically, alterations in the eye structure due to glaucoma occur before any noticeable functional effects. In order to detect glaucoma early, it’s important to classify these structural changes. The prime method for glaucoma detection includes examining the colour fundus images. This diagnostic tool can identify signs of damage to the ON associated with glaucoma like, an increased cup-to-disc ratio (CDR), rim thinning, notching, undermining, cupping, disc hemorrhage, and irregularities based on Retinal Nerve Fiber Layer (RNFL). Then after the effective method in characterizing glaucoma, both qualitatively and quantitatively, is optical coherence tomography (OCT) [9]. OCT primarily targets the optic nerve head and the macula region which providing high sensitivity and specificity in detecting preperimetric glaucoma [10]. Various OCT scan parameters, including disc topography and layer thickness measurements, effectively capture glaucomatous structural changes [11, 12]. This can advisable to utilize OCT information from overall the OD and macula as part of a thorough diagnostic strategy for identifying various types of glaucoma. Currently, significant advancements have been made in machine learning technologies, particularly in deep learning. This progress has facilitated the creation of novel models for automated diagnosis of eye diseases, like glaucoma. However, it is noteworthy that existing machine learning (ML) models in these studies focused solely on one catagory of image to segregation between glaucoma and healthy. This method significantly deviates from the thorough clinical examination carried out by ophthalmologists. Consequently, a limited number of ML techniques leverage fundus images relevant to glaucoma detection. ML method has been frequently employed to automatically identify different eye conditions, such as glaucoma[13], Diabetic Retinopathy (DR) [14], Age-related Macular Degeneration (AMD) [15], sevral retinal issues [16]. Currently, to demonstrate such Retinal Fndus Images (RFIs) which serve as a valuable tool in identifying numerous non-ocular conditions, including Type-II diabetes [17], anaemia [18], diabetes [19] and cardiovascular risks [20]. In the realm of glaucoma classification, different imaging modalities and clinical examinations are utilized, such as RFIs [21],OCT [22], and VFTs [23]. Despite the availability of multiple approaches, fundus imaging stands out as the most prevalent and cost-effective technique for widespread screening of diverse retinal diseases [24]. This study focuses on enhancing result optimization by combining multiple feature selection techniques with the ELM algorithm. The approach involves the utilization of IMGWO to achieve better classification accuracy. The study improves feature normalization and reduction by employing a hybrid method that combines Principal Component Analysis (PCA) and Linear Discriminant Analysis (LDA). Additionally, the study employs an IMGWO-ELM algorithm to classify glaucoma, improving the accuracy of the results.

Recently, the implementation of deep learning methods in the clinical setting remains constrained by certain limitations. The primary aspect is that deep learning approaches focus solely on input images and resulting outputs, acking clarity about how information moves through the internal layers of the network. In critical scenarios like fundus imaging, it’s vital to comprehend why the network makes its predictions, ensuring the model offers accurate estimations. As a result, there is significant interest in Explainable AI (XAI) to investigate the opaque DL models using the medical domain [25]. XAI techniques enable researchers, developers, and end-users to comprehend deep-learning models and explain their decisions to humans. Medical consumers increasingly demand explainability in deep learning to trust and implement these systems for biomedical domain [26]. Additionally, in GDPR, a law established by the European Union to regulate data protection, requires that automated learning systems offer an explanation prior to being used clinically with patients [27].

In this research, our objective is to create an automated XAI method based CAD model tailored for categorizing digital fundus images. The devised model incorporates an accelerated learning approach known called ELM [28, 29] to facilitate glaucoma classification. Furthermore, it integrates a customized iteration of improved grey wolf optimization (IMGWO) to enhance the model performance utilized on input weight and bias, thereby overcoming ELM’s inherent issues[30]. To elaborate further, the primary contributions encompass:

- Exploration of the FDCT-WRP technique to capture 2D singularities from glaucoma fundus images.
- The enhanced version of a single hidden layer feed-forward network (SLFN), referred from ELM, improves existing models for faster acquisition and enhanced generalization performance.
- Based on adopting a hybrid approach, incorporating improved grey wolf optimization and ELM, it aims to address challenges such as avoiding local minima, improving evaluating the speed of response in test data, and reducing the need for a huge number of hidden nodes based on the learning process.

The comprehensive documentation of this research endeavor is outlined below: the related works have been condensed in accordance with the specified Section 2. In Section 3 explained about the proposed methodology. It’s detailed simplification of the experimental result has been done in Section 4. Afterward, we share the ultimate conclusion and future work engage in a final discussion in Section 5.

## 2. Related works

In the field of medical imaging, explainable artificial intelligence (XAI) techniques are generally divided into two main groups: perturbation-based and gradient-based methods. Perturbation-based techniques involve analyzing the network by modifying input features and observing the effects of these changes on the model’s output predictions during the forward training phase. Some methods used in explainable artificial intelligence (XAI) are LIME, SHAP, deconvolution, and occlusion. These methods often rely on gradient-based approaches, which involve calculating the partial derivatives of the neural network’s output predictions with respect to input images [31]. These approaches are advantageous because they implemented after the training phase, avoiding based on trade-off between model accuracy and explainability. They are also typically faster than perturbation methods since their runtime hasn’t dependent based on list of input features. Back propagation-based techniques, like Vanilla Gradient, Guided Back propagation, Integrated Gradients, Guided Integrated Gradients, SmoothGrad, Grad-CAM, and Guided Grad-CAM are widely employed [32]. Although numerous XAI techniques have been devised for fundus images, there has less emphasis on elucidating brain imaging applications, especially concerning glaucoma detection. In one experiment, 2D Grad-CAM was employed to elucidate the workings of deep neural networks in identifying glaucoma. However, this method shares the limitations of previous classification explanation techniques, namely its restriction to 2D representations. Another approach, detailed in another study, aimed to overcome this limitation by extending class activation mapping (CAM) to generate 3D heat maps, providing a more comprehensive visualization of segmentation output. Despite its effectiveness in discerning classes, this method necessitated a balance between model complexity and transparency, thus impacting the effectiveness of ELM. Our recent years, a multitude of computer-aided diagnosis (CAD) schemes have emerged to classify glaucoma, emphasizing key elements like, pre-processing, extraction of features, feature dimensionality reduction, and classification. The aim is to gather and summarize the techniques and notable features of recent advancements in utilizing machine learning (ML) to categorize and diagnose glaucoma. Zhang et al. [33] introduced an innovative computer-Aided Diagnosis (CAD) model, which underwent experimentation to assess four ocular conditions. Shinde et al. [34], a novel computer-Aided Design (CAD) model incorporating contrast-limited adaptive histogram equalization (CLAHE) methods, have deployed on extraction of relevant feature from unlabeled datasets [30]. This approach has been implemented to mitigate the overfitting issue. Maheswari et al. [35] introduced an innovative method for glaucoma detection. Their approach involves employing Empirical Wavelet Transform (EWT) for image decomposition, extracting correntropy features, and employing the least-squares support vector machine (LS-SVM) used as glaucoma detection. Kansal et al. [36] have utilized characteristics derived from the dual-tree complex wavelet transform. This transform is used to apply fuzzy c-means clustering techniques and Otsu’s thresholding for segmenting the optic cup. In [37], the authors introduced a method for OD localization, employing the descriptor based on non-parametric GIST. This descriptor has applied on diminish the dimensionality using locality sensitivity discriminant analysis (LSDA) with several selection of feature and ranking techniques, followed by classification. Parashar et al. [38] used an innovative computer-aided design (CAD) method for diagnosing glaucoma, incorporating wavelet analysis to decompose fundus images into multiple modes. Following this, we obtained fractal dimension (FD) and various entropy measures to capture and build a least square SVM (LS-SVM) method based on several kernel functions. In [39], an innovative computer-aided design (CAD) model incorporating machine learning techniques is employed. They introduced a deep sparse auto encoder as part of this model, aiming to blend characteristics from deep and primary features. This design enhances the overall capability to represent advanced features and has the capacity to enhance the effectiveness of articulating high-level features. Additionally, the model incorporates L1 regularization to enhance the collaboration of deep features, particularly in scenarios where there is a need for more sample data. Contemporary literature underscores the notable importance of machine learning, especially within the domain of ensemble learning techniques. This proves especially advantageous in the biomedical field, even when datasets are limited. Currently, numerous models rely on machine learning approaches, yet prior research has yet to focus on ensemble methods for glaucoma classification. As a result, our proposed investigation centres on ensemble learning, harnessing the collective capabilities of XGBoost, SVM, and logistic regression (LR) to achieve better classification results in contrast to conventional models. In [40], the authors utilized discrete wavelet transform (DWT), histogram of oriented gradients (HOG) features and subsequently applied on ELM as a classifier. Additionally, in [41], an innovative Computer-Aided Design (CAD) model. This model involves selecting correlation attributes using a bio-inspired algorithm and the application of a KELM classifier that relies on salp-swarm optimization. In [42] the authors presented a novel method that incorporates speeded-up robust feature (SURF), histogram of oriented gradients (HOG) techniques utilized as feature extraction methods. From [43], the authors introduced a method for localized optic disc identification through bit plane analysis, utilizing wavelet feature extraction with optimized genetic feature selection. They incorporated various learning algorithms, ultimately employing SVM as the classifier. Kevin et al. [44], introduced computer-aided design (CAD) model utilizescharacteristics based on high-order cumulants (HOS) and applies Linear Discriminant Analysis (LDA) to diminish with less number of features. They utilized naive Bayes (NB) and SVM classifiers to identify and detect glaucoma. Rajendra et al. [45] introduced a glaucoma detection method that relies on extracted features, specifically the Gabor transform coefficients yield Kapoor entropies, kurtosis, energy, mean, Reyni, Shannon, and variance. The analysis involved ranking the features through the application of a t-test. In [46] the authors presented a new scheme using lib SVM, a sequential minimal optimization classifier using a wavelet-based feature extraction method for clinical implementation on glaucoma. In [47], the authors used a novel computer-aided design (CAD) approach has been developed for identifying glaucoma, relying on higher-order statistics (HOS) features, gradient information scale based on capturing the shape of features. PCA has been used for feature selection. In [48], the authors used a new model based on applied a Laplacian of Gaussian filter to isolate optical OD density in the red channel and multivariate with m-methods as classification. In [49], employed a CAD approch used on discriete transform(DT) and histogram equalization at pre-processing stage. They used DWT, HOS as feature extractors, SVM utilized on classification of glaucoma. In [50] presented a novel approch on hybrid optimization technique and hyper-analytic wavelet transformation method for extraction of features and SVM used for classification with radial basic function. Ananya et al. [51] suggested a novel CAD technique with HOG and FNN.

In literature, our main objective is to develop an improved GlaucoXAI framework provides 2D explainable sensitivity maps to help ophthalmologists comprehend and trust the performance of deep learning classifiers. we propose a method utilizing wavelet and its different forms, such as SWT, DWT, and FAWT, are commonly employed for extracting features. Hence, the conventional discrete wavelet transform (DWT) faces significant constraints, characterized by limited directional sensitivity and translation invariance, SWT has effectively address issues related to translation variance. Thus, it leads to repetition and fails to capture intricacies in high-dimensional singularities, rendering all these transforms less adept at addressing 2D singularities. Then, a combined reduction features is employed, known as the PCA+LDA method, is deployed on determine the most essential set of features. Ultimately, a refined training algorithm named IMGWO-ELM is presented for Single-Layer-FNN (SLFN). Such algorithm provides benefits such as evading local minima, enhanced generalization capacity, quicker learning pace, and improved conditioning matched with conventional algorithms namely FNN, SVM, LS-SVM, and ELM. So, to further enhance directional selectivity, it must be investigated to capture the detecting curve-like characteristics using images of the fundus. FNN and SVM have traditionally been observed to exhibit conventionally utilized even though they need list of parameters and more computational time. Hence, Many approaches have been tested on limited datasets and demonstrated elevated levels of accuracy; their performance is poor when the dataset is large. Hence, there is an opportunity to address the limitations of current approaches concerning the required number of features and enhance accuracy, mainly when dealing with extensive datasets. Here, the contribution based on threefold:

- An improved explainability framework, namely GlaucoXAI, has been employed in the recent ML models based on glaucoma detection making research interpretable without modifying the architecture.
- GlaucoXAI incorporated seven cutting-edge backpropagation XAI techniques to produce 2D visual representations of FDCT-WRP+PCA+LDA+IMGWO+ELM for glaucoma detection.
- An extensive assessment of the proposed framework demonstrated promising results in elucidating the outcomes for Glaucoma classification.

## 3. Proposed methodology

### 3.1. Proposed GlaucoXAI

In this paper, we have proposed an explainable artificial intelligence model named as GlaucoXAI (Glaucoma explainable artificial intelligence) The general flow of GlaucoXAI comprises into two primary components: a ML model specialized in processing fundus images, and a generator that provides explanations. We have consdered fundus images as input, then that images are forword propagated based on ELM classifier, which are then processed through task-specific computations to produce the desired output for a classification task. Later, The network’s results are displayed to medical professionals for assessment, with the possibility of asking for further explanation. Subsequently, Visual explanation maps are created by the explainability component to interpret the results of deep neural networks. Advanced explainable artificial intelligence (XAI) techniques can facilitate this process. Our GlaucoXAI framework extends deep image interpretation techniques, converting it into a multi-label detection task. It offers state-of-the-art XAI methods for both 2D and 3D medical image data. This research proposes a glaucoma diagnosis system that operates via the cloud and monitors health data from remote users to detect glaucoma diseases. The method can be easily adapted to diagnose and categorize different applications and determine whether an illness is glaucoma or healthy. The first step in this process is to use Principal Component Analysis (PCA) to identify important features, and filter out unnecessary ones. The proposed model is designed to initially learn how to reduce computational complexity, and then we have applied the Extreme Learning Machine (ELM) as a classifier. In our deployed scheme focused on four prime sections: preprocessing of images, extraction of features, reduction of feature dimension, classification. The glaucoma fundus image undergoes preprocessing, during which the area of interest (ROI) is isolated. This isolation relies on feature extraction utilizing FDCT-WRP. Then, prominent features have combined, deploying PCA with LDA techniques. Subsequently, The key characteristics are fed into the optimized IMGWO-ELM to facilitate classification. The detailed explanation of the deployed model’s framework and each of its sections is provided, shown in Figure 1.

**Figure 1.**
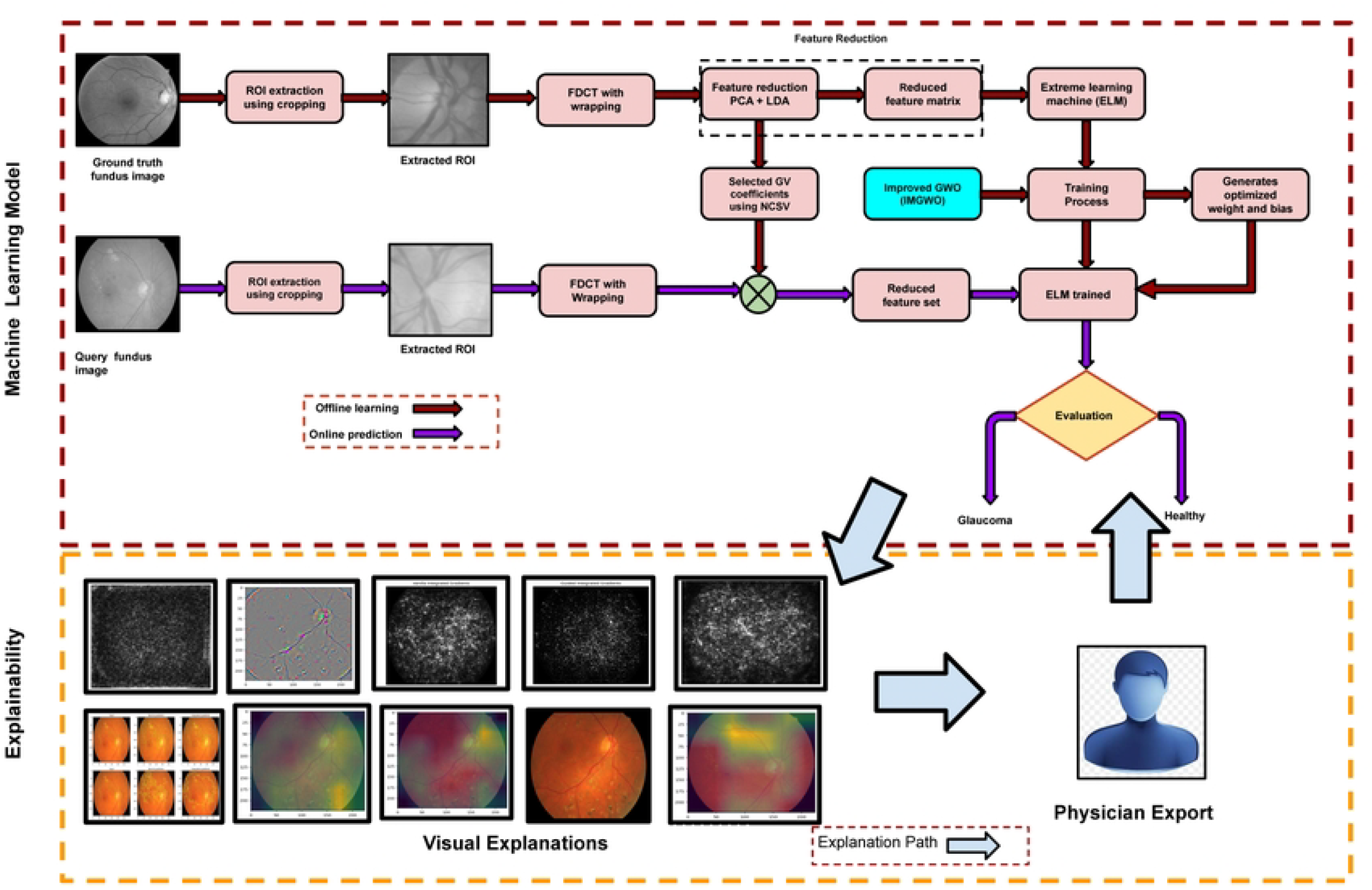
The sequence of the suggested GlaucoXAI framework.

### 3.2. Vanilla gradient

Vanilla gradient (VG) [52] represents the most basic method for visualizing areas within an image that have the greatest impact on the neural network’s classification outcome. It generates a saliency map by performing a single backward pass of the output class activation after completing a forward pass through the network In essence, it calculates the VG of the output activation concerning the input image. Suppose *P*_*c*_, is the prediction class c, evaluted by the classification of ELM for an input image *X*^*I*^. The prime aim of Vanilla gradient is to find the *L*_2_ regularized image, that has the maximum *P*_*c*_, while *λ* is the regularization term:

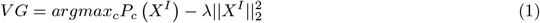

### 3.3. Guided backpropagation

Another approach to calculate the gradient of a particular output in relation to the input is to use guided backpropagation (GBP) [53]. The GBP introduces a novel variation of the deconvolution method aimed at highlighting the specific area in an image that triggers the highest activation for a particular class [54]. Let’s define F as the output from ELM, labeled as l and let B denote the resulting fudus image obtained through classification.

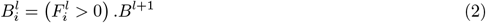

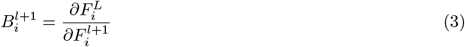

### 3.4. Integrated gradients

Sundararajan et al. [55] have proposed integrated gradients (IG) to address the saturation issue commonly encountered in gradient-based approaches. Suppose the function F : *R*^*n*^ → [0, 1] specified as deep neural network that has *X*^*l*^ = *γ*(*α* = 1) ∈ *R*^*n*^ as the input image, then *X*^*B*^ = *γ*(*α* = 0) ∈ *R*^*n*^ represented as baseline. The baseline is simply a black image with all values set to zeros. The IG can be evaluated using accumulating the gradients at all points on the straight-line path from the baseline *X*^*B*^ to the input fundus image as *X*^*I*^ :

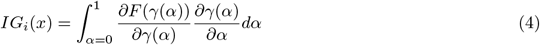

Here, i is the feature for the input image, whereas *α* shows interpolation constant to perturb image features.

### 3.5. Guided integrated gradients

Kapishnikov et al. [56] presented guided integrated gradints (GIG) as an aduption path based on the input image, baseline, and the deep model to be explained. Like IG, the GIG computes the gradients along the trajectory (c) originating from the baseline (*X*^*B*^) and concluding at the input under scrutiny (*X*^*I*^). Hence, The GIG path (c) is dynamically decided at each stage rather than adhering to the predetermined direction of the IG. Essentially, GIG identifies a subset of features (S) with minimal significance compared to all features in the input image. Mathematically,

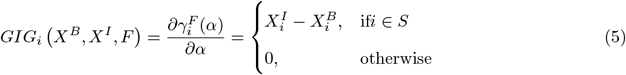

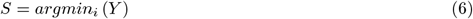

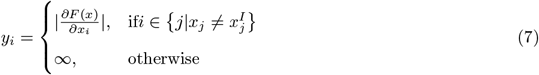

### 3.6. SmoothGrad

Smilkov et al. [57] smoothGrad introduced an improvement to address a prevalent problem in gradient-based techniques. SmoothGrad addressed this problem by generating visually enhanced sensitivity maps.It calculates the gradient across several samples around the input *X*^*i*^, and the average is determined after incorporating Gaussian noise.

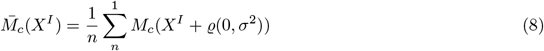

Here, *M*_*c*_(*X*^*I*^) specified as original sensitivity map, n is the number of samples, and *ϱ*(0, *σ*^2^) shows the Gaussian noise with variance *σ*^2^. Overall, *M*_*c*_(*X*^*I*^) shows any gradient-based visualization techniques, like explanation approches.

### 3.7. Grad-CAM

In [58] the authors introduced the class activation mapping (CAM) visualization techniques to a diverse range of ELM. The gradient CAM (GACM) used provides visual explanations without the need for retraining or enhancing the model architecture. Initially, the gradient for any target class *c* is calculated. Then, the activation feature map *M* of a specific layer *l* is globally averaged. First, the gradient for a particular target class *c* is averaged across all dimensions: width, height, and depth. The class-discrimination heatmap of GCAM is generated by applying a weighted combination of these activation maps, utilizing the ReLU function. Here, 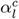 specified the nuron importance weights.

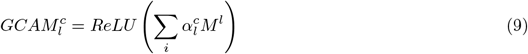

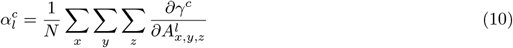

### 3.8. Guided Grad-CAM

GGCAM was introduced with the aim of offering enhanced visualizations at higher resolutions, effectively capturing intricate details of the subject in focus [58]. GGCAM combines the point-space gradient visualization technique GBP with the class-discriminative coarse heatmaps of GCAM by multiplying them element by element [53]. Before the point-wise multiplication with GBP, the saliency map of GCAM is upsampled to the input XI spatial resolution using bilinear interpolation.

### 3.9. Application to classification

In this paper, we present how GlaucoXAI can be utilized to produce visual interpretations for automatically grading brain gliomas using machine learning (ML). Our primary aim is to showcase the explanatory potential of our GlaucoXAI framework in aiding clinicians, rather than solely focusing on achieving optimal classification outcomes. Nonetheless, the classifier we employed demonstrated outstanding accuracy of –%, outperforming existing methods. To gain insight into the predictions made by the deep learning model, we utilized GlaucoXAI to produce a range of sensitivity maps, as depicted in Figure 2. These 3D visualizations of features were generated post-training. Explanation maps generated by methods (b-f) accentuate all influential features, whereas CAM heatmaps (g and h) emphasize the crucial regions of input images for distinguishing specific classes. Furthermore, visualization techniques such as GBP, IG, and GIG in pixel-space XAI methods emphasized intricate details within the MRI image but lacked distinctiveness in terms of class identification. Conversely, localization methods like GCAM produced highly distinctive activation maps corresponding to specific classes. Notably, combining GBP with GCAM resulted in superior localization with high-resolution visualizations. Smooth-Grad produced the most comprehensive feature maps, accentuating the primary discriminative regions in the FLAIR image for accurate glioma grading. In contrast, VG generated noisy visualization maps due to gradient saturation, rendering it less dependable for this application compared to other methods, as noted in [59].

**Figure 2.**
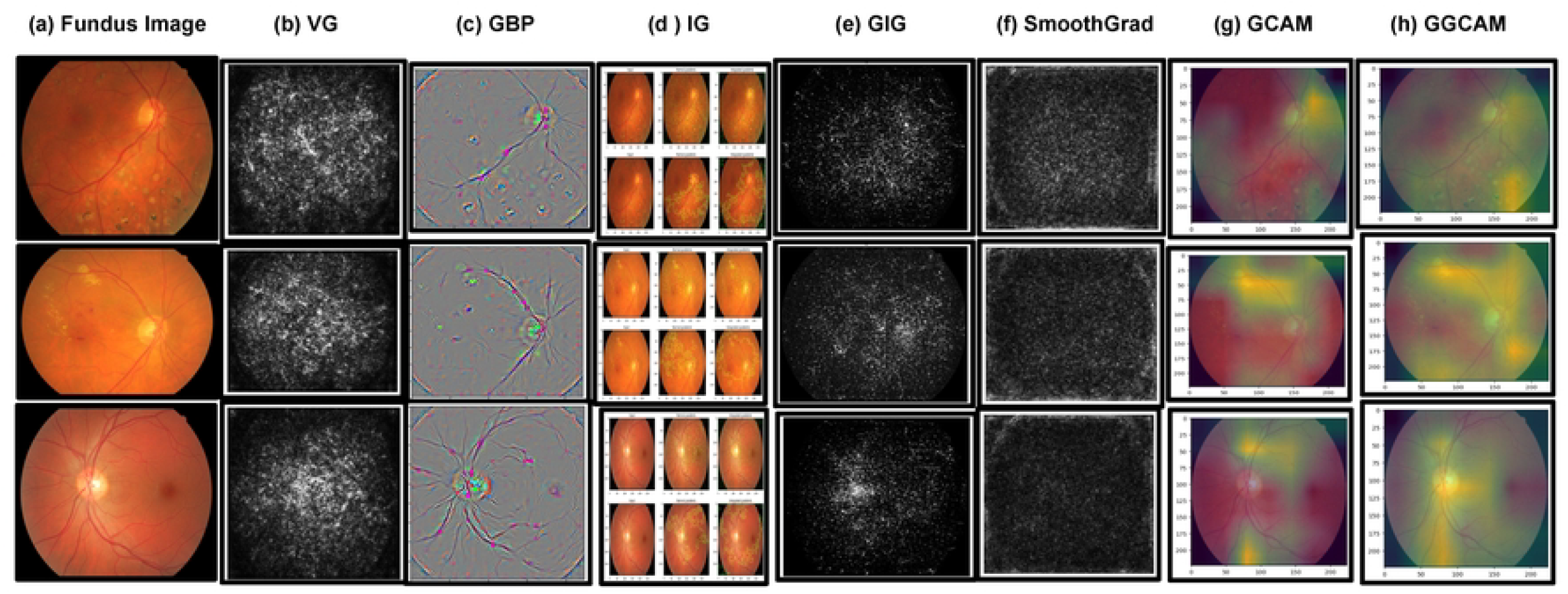
Comparing different explainable artificial intelligence (XAI) visualization methods for the classification of glaucoma. Sensitivity maps are presented for High-Grade Glioma (HGG) cases in the top four rows and for Low-Grade Glioma (LGG) cases in the bottom three rows. The arrangement displays various visualization techniques applied to original fundus images, progressing from left to right: Vanilla gradient, guided backpropagation, integrated gradients, guided integrated gradients, SmoothGrad, Grad-CAM, and guided Grad-CAM. It’s observed that in certain techniques (b, c, d, e, f), all relevant features are highlighted in white, whereas in others (g, h), red regions indicate a high score for the predicted class

#### 3.9.1. Preprocessing of images

During preprocessing, fundus images consists of several undesirable elements namely, noise, artefacts, test samples, etc., at the time of acquisition. List of anomalies continue to spread to the extracted features unless eliminated beforehand. Therefore, conventional image preprocessing approaches, such as increasing noise reduction techniques, have been applied. To expedite processing, regions of interest (ROIs) are manually extracted using cropping methods [60, 61] for two standard datasets, namely G1020 and ORIGA based on x, y and centre co-ordinate values specified through ophthalmologists. In Figure 4 display selected sample and extracted Region of Interest (ROI) images. Our proposed approach involves utilizing cropped images sized at 128×128. In contrast to relying solely on limited knowledge from specific ROIs, our method considers the entire image, also sized at 128×128.

#### 3.9.2. Feature Extraction using FDCT-WRP

The utilization of wavelet transform is widespread owing to its multiresolution characteristics and the capability to provide time–frequency localization information for an image. Nevertheless, conventional wavelet methods cannot detect two-dimensional singularities such as lines and curves, in contrast to ridgelet and curvelet transforms [62]. The Ridgelet transform can effectively capture diverse characteristics such as straight lines and arbitrary orientations; however, it needs to be better suited for handling curve singularities found in an image[63]. The issue is addressed by a fast discrete curvelet transform, which utilizes on multi-scale ridgelet specifed [64] and presents various features, including enhanced directional selectivity, multiresolution capability, anisotropy, and localization. The curvelet is an alternative form, known as curvelet transform of second-generation, was introduced [65]. It reduced the limitation of under-recognition of ridglets and computational overhead based on first-order curvelet transforms specified [66]. The prime derivation in mathematics is the discrete curvelet transform is explained as follows:

Based on signal *S*_*i*_, the transform of curvelet *CU*_*r*_(*p, q, r*) has been specified on the dot product of *S*_*i*_, Ψ_(_*p, q, r*) which is

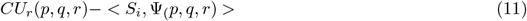

Here, Ψ_(_*p, q, r*) denoted as a curvelet basic method, and *p, q, r* has denoted as a number of parameters, namely scale, position and orientation accordingly. The technique of curvelet transformation entails the segmentation of each image based on multiple windows across several scales and orientations. Based on depiction on a CT through an input fundus image in a discrete form is *f* [*a*_1_, *b*_1_] with 0 ≤ *x*_1_, *y*_1_ *< m* is shown in :

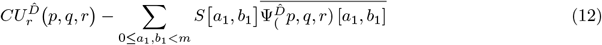

Here, 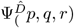 shows a DWT. Then, the CT of the second generation has the capability to generated using two distinct approaches known as wrapping with Unequally Spaced Fast Fourier Transform (USFFT). Unlike, the first-generation curvelets, that methods have known for their simplicity, speed, and decreased redundancy. However, when comparing the two techniques, it is evident that FDCT-WRP employing the method of wrapping which notably easy to use, more straightforward for utilization, faster than those using USFFT. Recognizing these advantages, we have opted for the wrapping schemes to develop referred to as FDCT-WRP, serving as extraction of features. The sequence of instructions for building that is FDCT-WRP is outlined below:

Evaluate the fast Fourier transform (FFT) based on the two-dimensional coefficients (*V*_*e*_ [*c*_1_, *d*_1_]) for each image.

Create the discrete localization window that corresponds to each scale and angle within the fourier domain 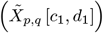 and evaluate the product 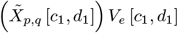

By circularly wrapping the data around the origin and re-indexing it 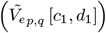

To acquire the DCT coefficients 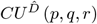, applying then 2D FFT inverse using 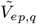

Here, a feature vector is formed by attaching the FDCT coefficients through a wrapping techniques at every scale based orientation, similar to how p, q are employed. For calculate the list of scales based on size of image *m*_*r*_×, *m*_*c*_ as follows:

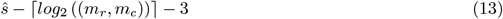

Each image has size 128 × 128, the *ŝ* denoted as 4. It implies that every image undergoes decomposition into four levels in the curvelet transform. Scales 2 and 3 encompass distinct sub-band information except for the initial and final scales. At angles *α, α* + *π*, as curvelet generates identical coefficients. Therefore, we have eliminate a half-coefficient based on sub-band at every scale. Therefore, The values of the feature vector’s coefficients are notably elevated, which is crucial to further reduce them in order to isolate the promenent features.

#### 3.9.3. Dimensionality reduction of Feature vector

Reducing the dimensionality of features plays a vital role in predictive modelling and tasks related to machine learning. Storing and processing feature vectors with a higher dimension requires increased storage capacity and more significant classification computational resources. Here, commonly utilized method named as PCA, has employed in initial stage. In the subsequent phase, linear discriminant analysis (LDA), a supervised technique capable of identifying pertinent features and distinguishing them within comparable classes, is utilized to maximize benefits [67, 68]. The features acquired through PCA have been leveraged by LDA for further dimensionality reduction. Firstly, The 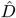 denoted as dimensional reduction feature that reduced through the application of PCA to decrease it to M dimensions. After that, LDA was used to further reduce the dimensionality to L, resulting in the final reduced dimension is L ¡ M ¡ 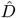 [69, 60, 62]. Sorting the eigenvalues of the feature list in descending order is undertaken to pinpoint the most optimal features. Then, the application of a normalized cumulative sum of variance (NCSV) technique was initiated for assess the NCSV values for each feature. By using *i*^*th*^ features of NCSV values are computed as,

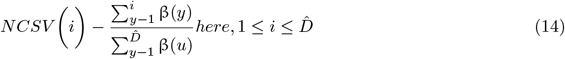

Hence, β(*y*) denoted as the eigenvalue of *y*^*th*^ feature, The dimensionality of the feature vector is 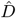. Here, the threshold has utilized for choose *L* as list of eigenvectors values based on NCSV.

### 3.10. IMGWO-ELM

This section describes Grey Wolf Optimization (GWO) and Extreme Learning Machine (ELM), and then introduces the proposed IMGWO-ELM algorithm.

#### 3.10.1 Grey Wolf Optimization

Optimization algorithms are pivotal in training machine learning models, parameter tuning, operations research, and solving real-world problems where finding the optimal solution is critical. The selection of the algorithm frequently relies on the particular attributes of the given problem, such as the nature of the objective method, presence of constraints, and the computational resources available. There is a list of optimization methods like Gradient Descent, Genetic Algorithms(GA), Simulated Annealing(SA), Ant Colony Optimization (ACO), Particle Swarm Optimization(PSO), Linear Programming, quasi-newton methods, and Gayesian Optimization. GWO is a swamp intelligence method like ACO proposed by Mirjalili [70]. Its outcome is better and the best optimization technique than the genetic algorithm. It primarily mimics the leadership hierarchy of wolves. It is simulated by grouping the population of search agents into four types of individuals according to their fitness, namely α, β, Δ, and ω. Here, α is denoted as the best-fit solution and ω is specified as a solution of least-fit.

Then, α, β, and ΔGuide the ω wolves to navigate the search environment for the prey, as described in Yang et al. [67]. The wolves update their position based on their encircle of the prey, which is followed by a mathematical Equation:

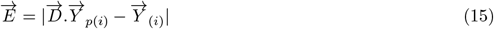

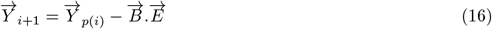

Here, *i* denoted as the current iteration, 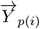 viewed as the current position of the prey, and 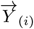 shows the current position of the wolf. The vector 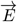 represents as the distance between the wolf and its prey. Additionally, the vectors 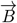 and 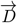 are defined as:

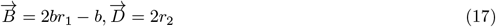

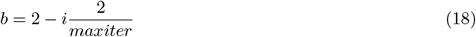

Now, 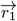 and 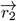 having two discrete vectors with whole values in 0 to 1, *i* specified as recent iteration. Finally, to achieve the best three solutions, ensure your information accurately reflects the current state of the prey; accordingly then the best three wolves wolves have been chosen by α, β, δ [71]. Finally, the remaining wolves, having ω, modify their placement in relation to the optimal choice among the three. The redistribution of wolves is implemented according to the subsequent equation:

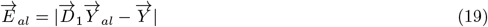

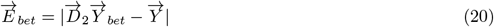

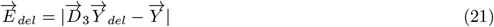

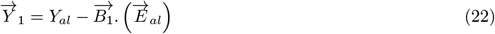

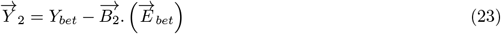

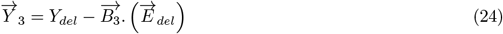

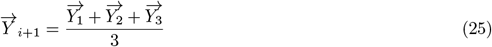

Here, 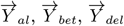 specifed as the location based on α, β, δ wolves, 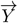 shows the present position of the solution *D*_1_, *D*_2_, *D*_3_, *B*_1_, *B*_2_, *B*_3_ are three randomly generated values. nomous and random allocation of hidden node parameters [72, 73].

#### 3.10.2. Extreme Learning Machine

ELM provides a simplified, streamlined, and effective approach for training neural networks, making them suitable for specific applications prioritizing quick training and simplicity. ELM has been utilized in a range of domains including pattern recognition, image and signal processing, and regression tasks. In contrast to conventional feedforward neural networks (FFNNs) that rely on the iterative assessment of all network parameters, the ELM approach is introduced, emphasizing the use of autoIt to address regression and classification challenges. Unlike standard neural networks, Extreme Learning Machines (ELM) employ a fixed assignment of weights and biases in their hidden layers, eliminating the necessity for the iterative training process typically associated with gradient descent methods. The core idea of ELM is to randomly assign weights and biases in the hidden layer based on the input. Then, it determines the output layer weights through an analytical solution based on a linear system solver. This approach enables ELM to avoid the lengthy backpropagation process, leading to significantly quicker training durations compared to conventional neural networks. During training phase of ELM, the network takes input data and computes the activations of the hidden layer by applying a non-linear activation function to the weighted sum of the inputs. The weights of the output layer are determined by solving using techniques like the Moore-Penrose pseudoinverse or other methods of regularization. Once training is finished, the ELM model applies the acquired weights to input features to forecast new, unseen data. Its appeal has grown due to its computational efficiency, particularly beneficial for handling extensive datasets.

Here, *Sa* specified as random distint samples (*p*_*i*_, *q*_*i*_), here *p*_*i*_ = [*p*_*i*1_, *p*_*i*2_, *p*_*i*3_, …, *p*_*in*_]^*T*^ and *q*_*i*_ = [*q*_*i*1_, *q*_*i*2_, *q*_*i*3_, …, *q*_*im*_]^*T*^ *ϵ* ℜ^*m*^. A conventional single-layer FFN (SLFN) with ’*H*’ hidden nodes specified by mathematically which follows:

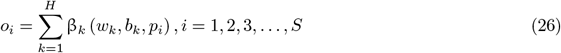

In this context, *w*_*k*_ and *b*_*k*_ represent the parameters associated with the hidden node, encompassing its weight and bias values. The vector *b*_*k*_ = [β_*k*1_, β_*k*2_, …, β_*km*_]^*T*^ serves as a representation denoting the output weight from the *k*^*th*^ hidden node to the output nodes. The result linked to the *k*^*th*^ node, denoted as *A*(*w*_*k*_, *b*_*k*_, *p*_*i*_), is equivalent to *p*_*i*_, where *o*_*i*_ represents the true output related to *p*_*i*_. Additionally, the mathematical representation of *A*(*w*_*k*_, *b*_*k*_, *p*_*i*_) have been expressed by:

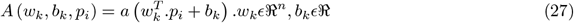

Here, a(*p*) : ℜ → ℜ specified as the sigmoid function 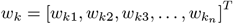 considered as the weight vector based on input, *k*^*th*^ hidden nodes, and *b*_*k*_ identified as the bias of the *k*^*th*^ nodes. The SLFN can be assessed for *H* hidden nodes across *S* samples without any error. In this context, then the cost function 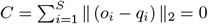. However, there exist (*w*_*k*_, *b*_*k*_) and β_*k*_ like,

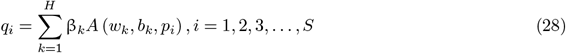

In Equation 28, the specification is presented concisely as follows:

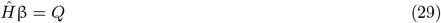

Here,

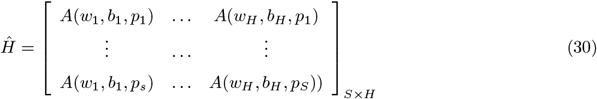

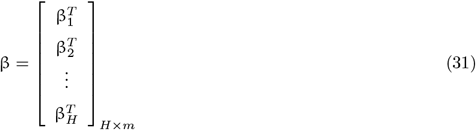

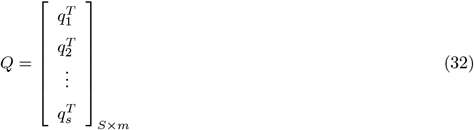

In this context, 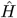 is defined as the matrixdepicting the outcomes of the hidden layer in the SLFN. In Equation 29, formulated based on linear method according to [64], characterizes the output weights associated with the model as:

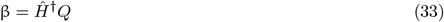

Here, 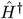 specified on the Moore-Penrose (MP) specified as the opposit of 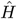 [65]. Hence, the researcher is specified as MP specification inverse explained in [66] for better clarity. The algorithm explained its stages in the ELM method viewed as:

##### Algorithm 1

Extreme learning machine (ELM) Algorithm pseudocode

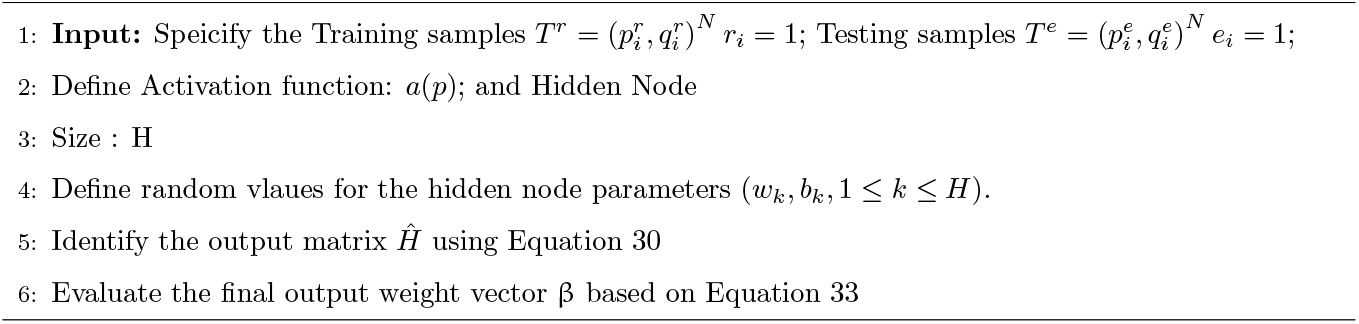

#### 3.10.3. Improved GWO-based ELM

From Section 3.10.2, we have noticed that, the ELM effectively applies its generalization characteristic to two specific parameters, namely, the weights and bias. Hence, the random initialization of hidden node parameters in ELM introduces two primary challenges, which can be readily observed [74]. Firstly, ELM exposes a significantly larger quantity of concealed neurons in contrast to conventional gradient-based algorithms. As indicated by the performance results of ELM, its speed diminishes when introduced with new test samples. Secondly, ELM focused on ill-conditioned hidden layers, resulting in vectors that evaluate the least generalization outcomes.

To address this problem, several evoltionary and swarm intelligence-based techniques have been examined and utilized [75, 76, 71]. These algorithms are selected because of their superior capability for conducting comprehensive searches on a global scale to address optimization challenges. In[75] presented a novel modified algorithm like in the Evolutionary-ELM (E-ELM), an improved version of differential evolution (DE) has been utilized to optimized the hidden parameters of ELM, specifically weights & bias. The solution is obtained through inverse utilization of MP generalization. Several researchers demonstrated as E-ELM exhibits superior, quicker effectiveness than traditional algorithms. Nonetheless, a drawback of E-ELM is the necessity to fine-tune two extra parameters, precisely, mutation and crossover. Xu and Shu [76] proposed an approach that combines PSO with an ELM for optimize it’s parameters of ELM. Furthermore, they established a specified range within PSO to increase it’s overall efficacy of ELM. Han et al. [71], have enhanced version of the ELM, referred to IPSO-ELM, has presented as attain optimized SLFNs using a PSO approach.

The effectiveness of the mentioned methods is reliant on the specific parameters of the algorithm, despite ongoing efforts to optimize these parameters. Therefore, establishing appropriate values for these parameters is critical on any domain of difficulty. Additionally, nadequate tuning of parameters may lead to notable computational complexity or becoming stuck in local optima [28]. In the case of Differential Evolution (DE), two essential parameters are essential: the scaling coefficient and crossover probability. Likewise, Particle Swarm Optimization (PSO) relies on the inertia weight and acceleration factor. This research employs a modern global optimization algorithm without the need for explicit parameter settings, specifically the GWO, to address the challenges associated with improper parameter tuning. Furthermore, a novel approach, named

IGWO-ELM, is introduced, merging the benefits of Grey Wolf Optimization with ELM.

#### 3.10.4 Improve the Convergence Factor

The convergence factor emphasizes the search ability of GWO. Linear convergence is ineffective for achieving a thorough global search and tends to get trapped in local searches quickly. Usually, in global optimization algorithms, the initial phase involves exploring a large search space, which takes more time to find the optimal value. After undergoing several iterations of improvement, the algorithm typically achieves the best solution faster. Because of its similar traits to the behavior mentioned earlier, the adaptive non-linear convergence factor linked with the exponential function intensifies this convergence phenomenon even more. The non-linear decline in value occurs as increased in list of iterations, exhibiting a gradual decrease in the early stages with a minor contraction range. This enables a meticulous fine search. Subsequently, in the later stages of the iteration, the value undergoes rapid changes, facilitating an expedited search process. Using the Grey Wolf Optimizer (GWO) to dynamically find the best parameter values for the hidden nodes of the Extreme Learning Machine (ELM), and subsequently applying the Moore-Penrose pseudo-inverse for obtaining an analytical solution. Additionally, it is essential to note that the Improved GWO (IGWO) algorithm incorporates an exponential function, resulting in a modification of Equation 18. Therefore, the exponential function exhibits symmetric characteristics, demonstrating an adaptive nonlinear convergence factor. in Equation 34 has been employed, that can achieve a variable convergence rate that adjusts nonlinearly [0,2] based on original linear convergence factor. To visually grasp the variation of the value ‘a’ across multiple iterations, refer to Figure 3. is graphically represented.

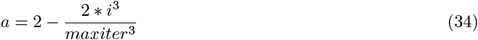

**Figure 3.**
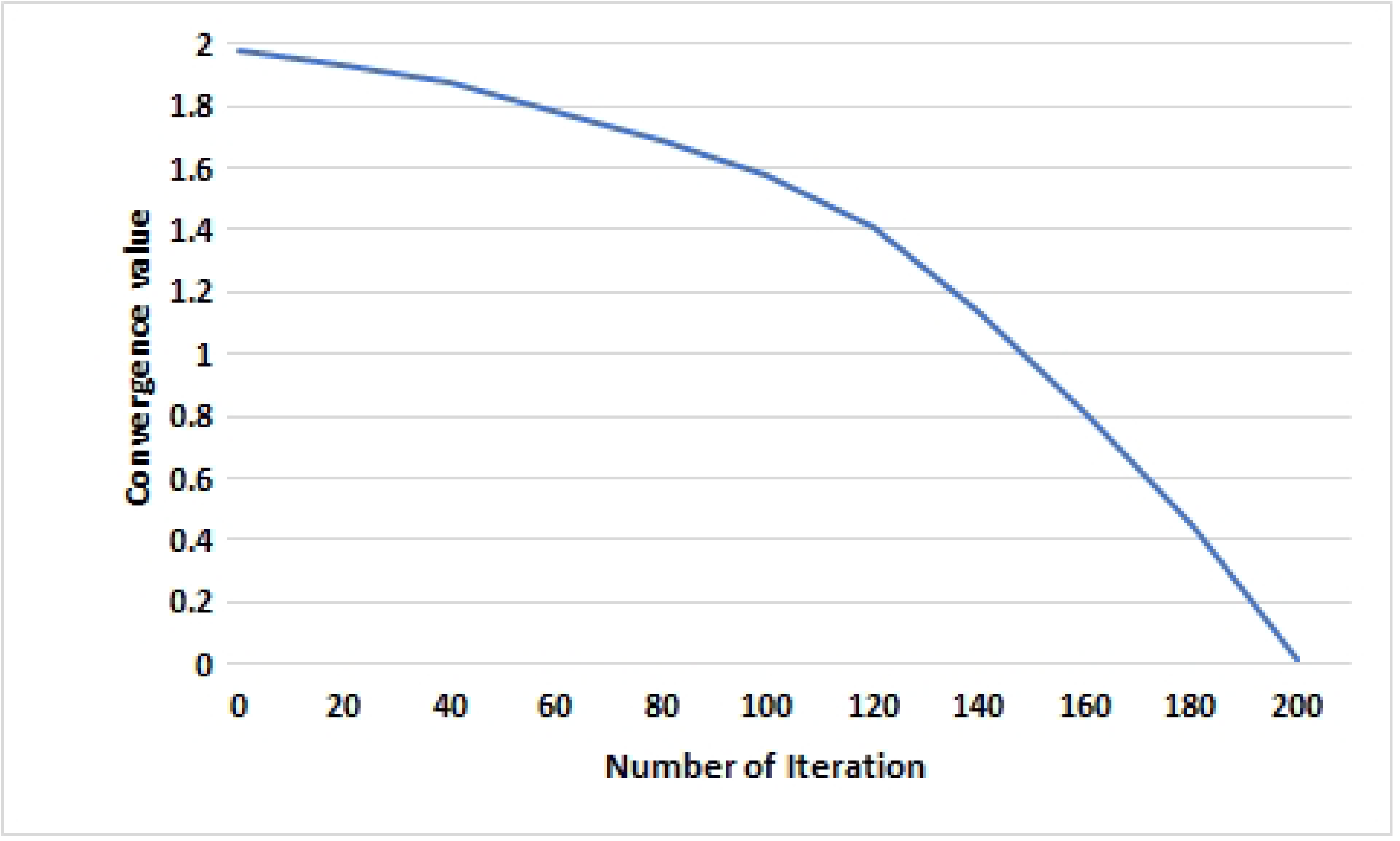
Graph of the convergence factor of the proposed model

**Figure 4.**
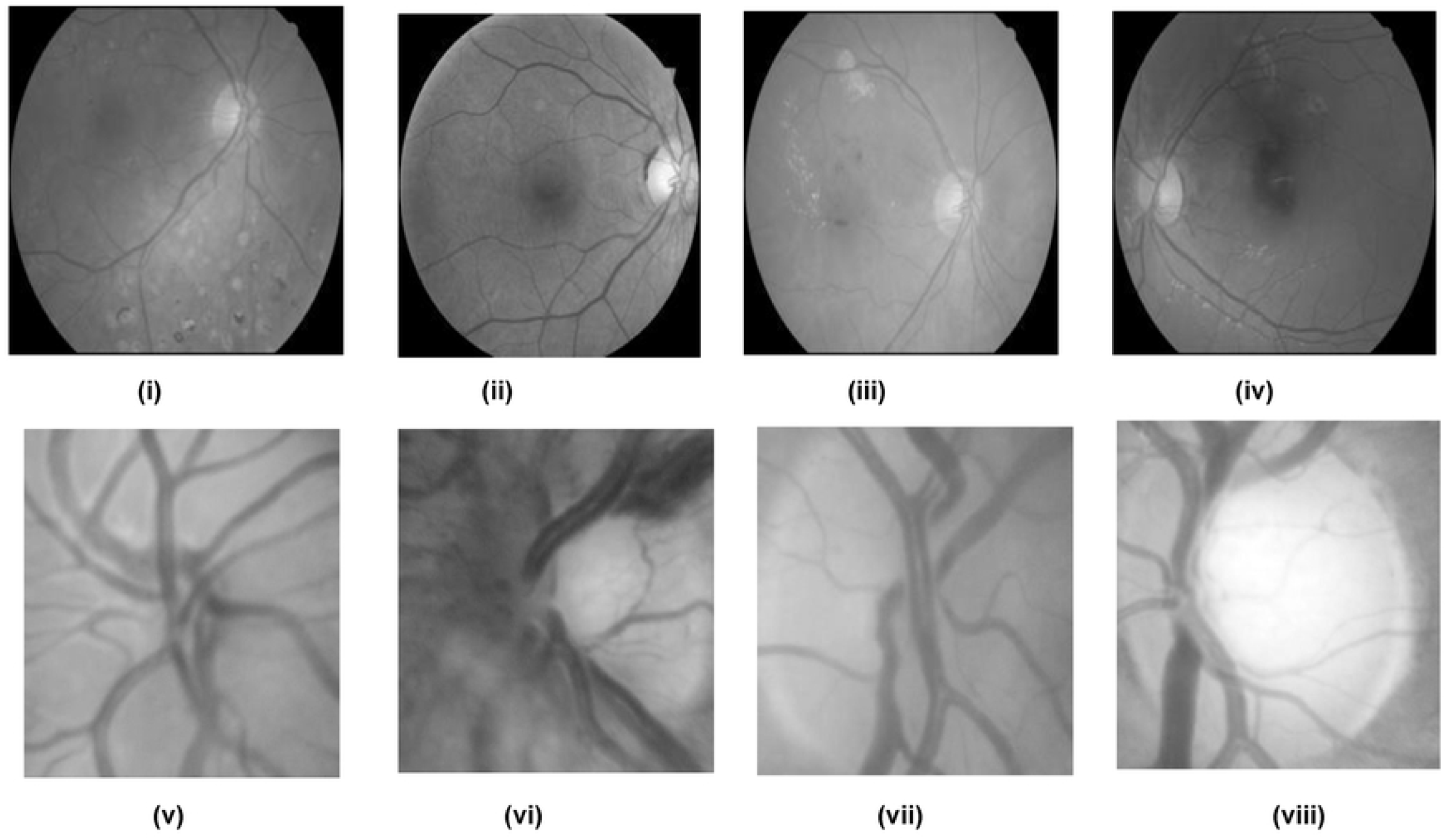
Some sample images of G1020 and ORIGA Dataset.

The IMGWO+ELM system consists primarily of two components: one focused on optimizing parameters and the other on evaluating classification. The primary objective of IGWO is to enhance classification accuracy while ensuring that input weights, hidden bias remain based on specified range. This is done for enhance the convergence characteristics of the ELM. The subsequent passage provides a brief overview of the various stages involved in IMGWO-ELM.

- Initially by randomly initializing each solution within the population of candidate. Each resolution comprises a collection of input weights and a bias organized in the subsequent manner:

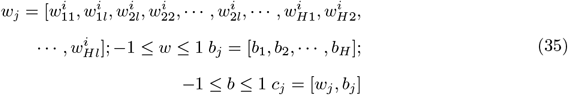
- Assess the classification accuracy and output weights for each potential solution. Specifically, evaluate the fitness value, which represents classification accuracy, based on the validation set to prevent overfitting concerns.
- Determine the three most effective outcomes (α, β, δ). Revise the outcomes by making updates in Equation 26.
- Generate innovative solutions by employing the fitness value in the following manner:

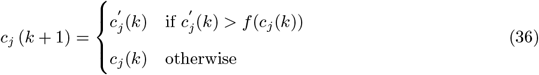 During this context, *f* (*c*_*j*_(*k*)) and 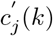 denote as the fitness of the *j*^*th*^ candidate solution and its corresponding updated solution based on *k*^*th*^ iteration, respectively.
- Search for occurrences of cases beyond bounds in each of the recently created solutions and limit them to a range of [-1, 1] using the following method:

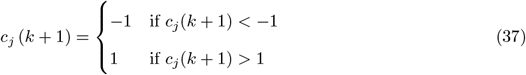
- Continue iterating through steps (III) to (V) until the termination state is reached. The termination state is the target fitness value to be achieved within the specified number of iterations. Afterward, the optimized parameters of the hidden nodes are essential and are used on the testing sample to assess the overall performance of the implemented scheme.

The algorithm 2 outlining the deployed scheme is presented in the pseudo-code

##### Algorithm 2

The Proposed model (FDCT-WRP+PCA+LDA+IMGWO) Algorithm pseudocode

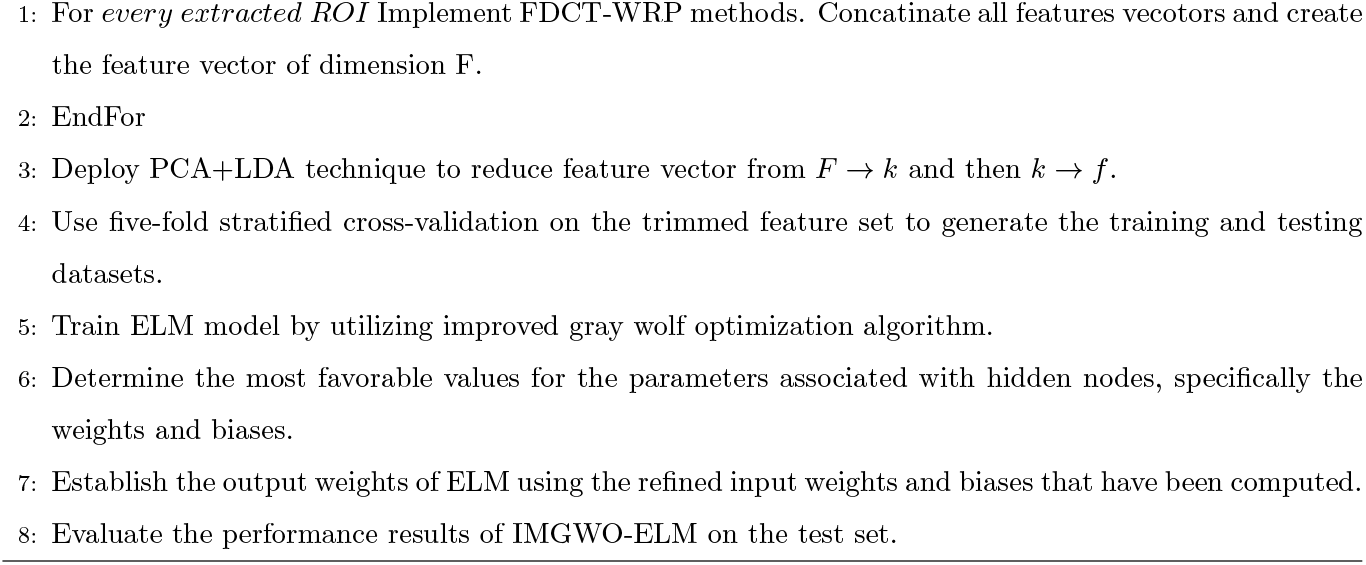

## 4. Experimental Result and Discussion

All experiments were conducted using the proposed CAD model in MATLAB R2018a on the PARAM Shavak–Supercomputer incorporates an HPC system and is embodied in a tabletop system that integrates an Intel(R) Xeon(R) Gold 5220R CPU @ 2.20GHz. The setup consists of a minimum of two multicore CPUs, each featuring a minimum of 12 cores. Furthermore, it is equipped with either one or two NVIDIA K40 accelerator cards and NVIDIA P5000 many-core GPU accelerator cards. This advanced system achieves a maximum computing capability of 3 Tera-Flops, supported by an 8 TB storage capacity and a substantial 64 GB of RAM. Additionally, it comes with a pre-installed parallel programming development environment, providing computing power equal to or exceeding 2 TF. During the experiment, We used two frequently employed datasets, specifically G1020 [77] and ORIGA [78], accordingly. We resized the image dimensions of 128 × 128 to ensure consistency.

- True Positive Rate (Sensitivity): It indicates the ratio of correct predictions for the positive class. 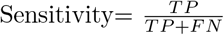
- True negative rate (Specificity): The probability that a true negative will lead to a negative test result. 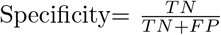
- Accuracy: It is characterized as the count of accurate predictions out of the overall predictions made.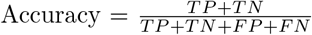 Where,
- TP (True positive)-Accurately identified positive instances
- TN (True Negative)-Accurately identified instances of negative cases.
- FN (False Negative)-Positively identified cases that were classified incorrectly.
- FP (False Positive)-Negatively classified cases Incorrectly.

### 4.0.1. Dataset used

The utilized approach has been verified using two commonly accepted datasets that consist of glaucoma fundus images, specifically G1020 [77]and ORIGA [78]. The G1020 dataset comprises a substantial set of retinal fundus images, openly accessible to the public. It is specifically created for glaucoma classification and consists of 1020 fundus images, including 724 depicting healthy conditions and 296 showing cases of glaucoma. Likewise, The ORIGA dataset, a widely recognized and extensively utilized resource, plays a prominent role in glaucoma research and analysis.This dataset includes 650 fundus images, with 482 representing individuals in good health and 168 featuring patients diagnosed with glaucoma. An elucidation of both datasets is provided in Table 1. Illustrations of representative images from both datasets are depicted in Figure 4. The distribution of trial samples through five-Fold stratified cross-validation in each iteration shown in Figure 5.

**Table 1:** Fundus images of Glaucoma Data Sets (Glaucomatous vs. Healthy)

| Data sets | Total images |  | Training images |  | Testing images |  |
| --- | --- | --- | --- | --- | --- | --- |
| | $G_l$ | $H_e$ | $G_l$ | $H_e$ | $G_l$ | $H_e$ |
| **G1020 [77]** | 296 | 724 | 178 | 434 | 118 | 290 |
| **ORIGA [78]** | 168 | 482 | 101 | 289 | 67 | 193 |
| $G_l$-Glaucoma, $H_e$-Healthy | | | | | | |

**Figure 5.**
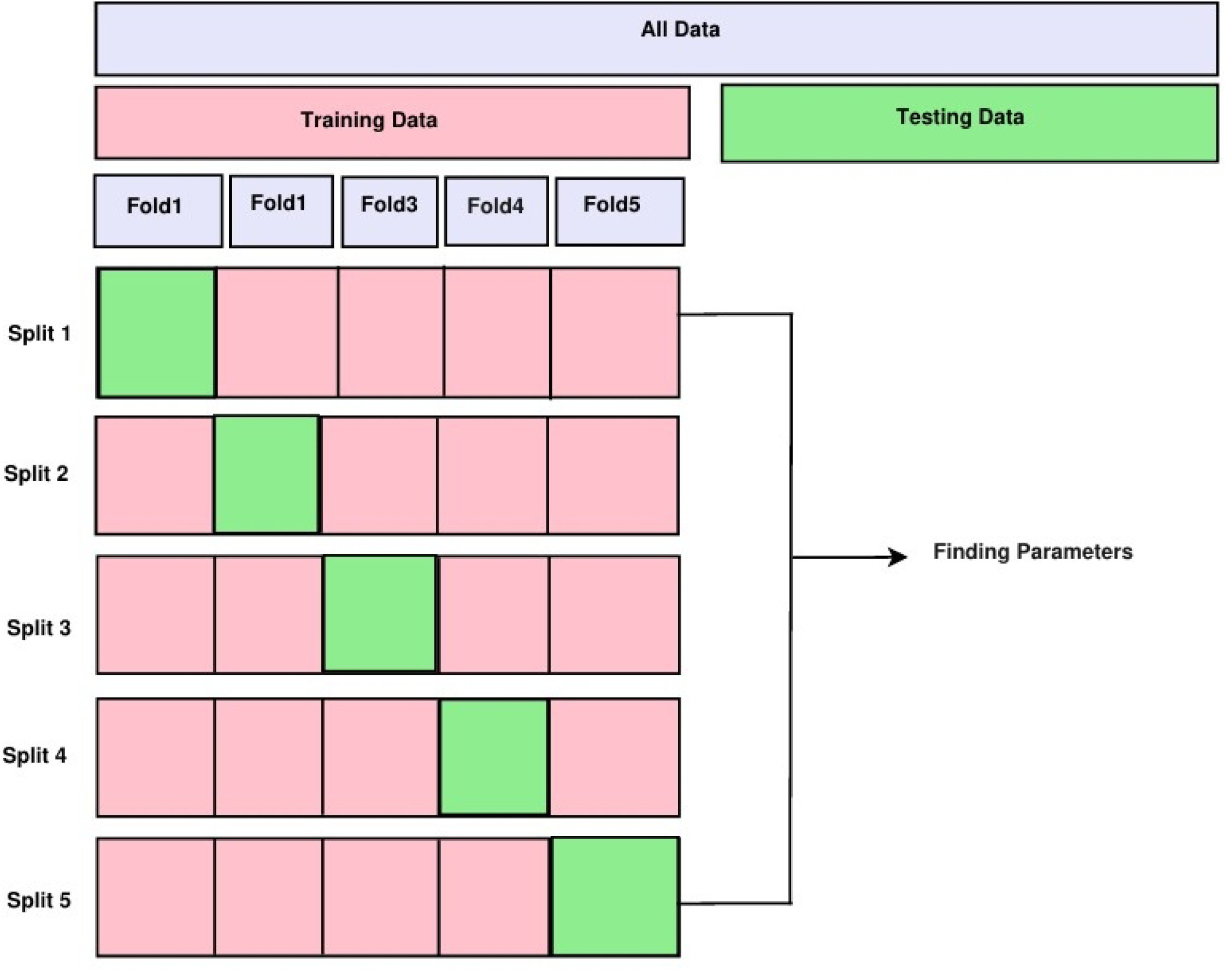
The distribution of trial samples through k-Fold Cross-Validation in each iteration.

### 4.0.2. Results of feature extraction and reduction

From our literary work, we incorporated the FDCT-WRP as a method for extracting characteristics. Following the application of decomposition using the specified Equation 14., each image is divided into four levels. After undergoing the decomposition at the fourth level, Figure 6 visualizes the creation of 50 sub-bands in total. The glaucoma detection based on digital fundus images involves scrutinizing the coefficients derived from the 4^*th*^-level curvelet decomposition. To streamline the feature vector, 25 sub-bands have been chosen from 50, excluding equivalent sub-bands. However, the range of features linked to an individual glaucoma image is extensive, encompassing 42,257 elements, a notably sizable quantity. To select the most relevant features, a combined approach known as PCA with LDA utilized to decrease feature dimensionality. The aim was accomplished by employing the normalized cumulative sum variance (NCSV) metric, that sought to identify and select relevant features from an initial pool of 42,257 features. The NCSV values of numerous features were individually evaluated for both PCA and the combination of PCA and LDA. Through experimental observation, it was determined that the PCA combined with the LDA approach incorporated additional information, utilizing 33 features and outperforming PCA alone. Here, we have manually assigned a value of 0.97 to NCSV. Consequently, our investigation revealed that the employed PCA-LDA technique achieved higher accuracy by utilizing 27 features in both datasets, surpassing the performance of PCA alone. Consequently, the same features were applied to the G1020 and ORIGA datasets.

**Figure 6.**
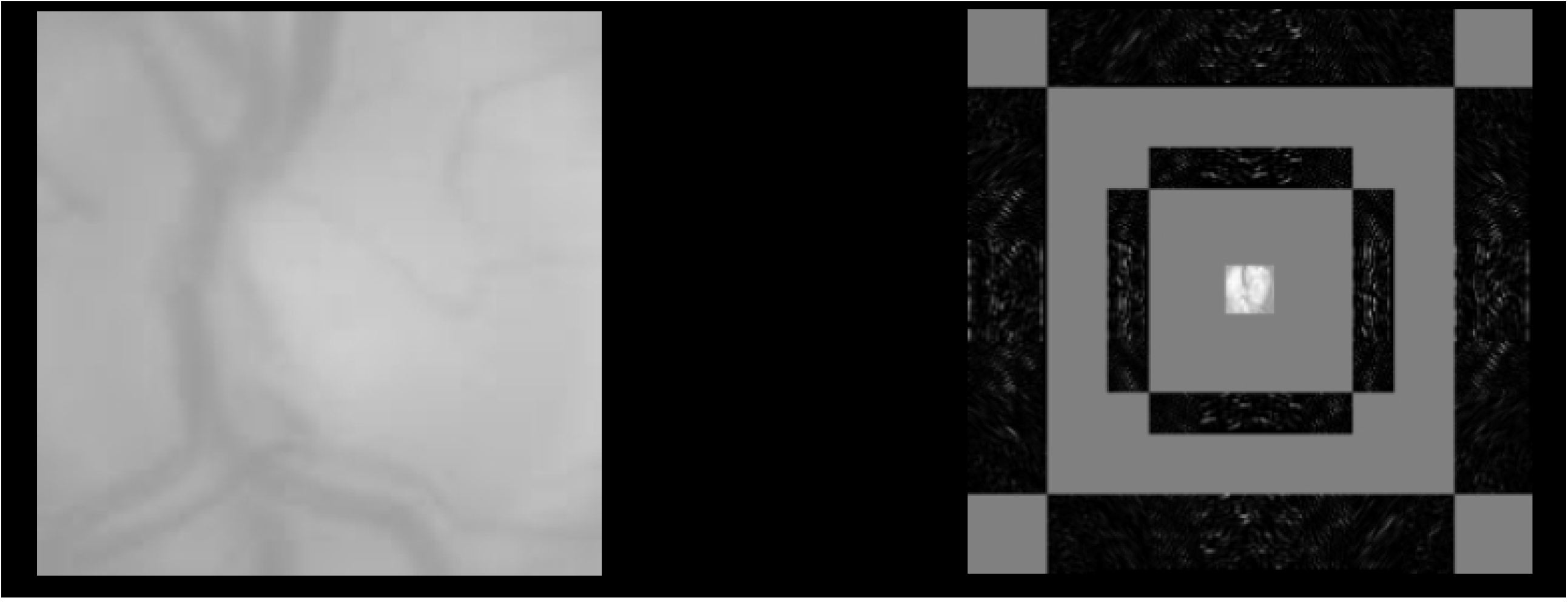
Glaucoma fundus image and its FDCT scale four coefficents

### 4.0.3. Classification Results

The deployed work has been implemented using an upgraded iteration of the CAD model called FCDT-WRP+PCA+LDA+IMGWO+ELM, designed for glaucoma detection. We have utilized the suggested model outlined, employing three distinct outcome matrices: accuracy, sensitivity, and specificity. The IM-FGWO+ELM is evaluated against various learning classifiers such as KNN, SVM, BPNN, and ELM. Table 2 illustrates the experimental parameters utilized in the IMFGWO+ELM model and various classifiers. Specific values, denoted as *c*1 and *c*2, have been assigned to the coefficients of both methods, GWO+ELM. Our IMGWO+ELM approach achieves superior outcomes with fewer hidden nodes compared to list of algorithms namely, KNN, SVM, BPNN, ELM, GWO+ELM, and IMGWO+ELM on two standard datasets. Additionally, our method shows lower condition numbers and norm values, resulting in improved classification effectiveness. In this study, we replicated the uniqueness of an operational model using twenty-seven features. The employed scheme has assessed against state-of-the-art models from diverse classifiers. Likewise, In the G1020 dataset, accuracy is exhibited by algoriths like, KNN, SVM, BPNN, ELM, MFO-ELM, GWO-ELM, IMGWO+ELM, namely, 90.69%, 91.67%, 92.65%, 93.63%, 91.25%, 93.40% and 93.87% accordingly. Similarly to the ORIGA dataset, the list of models like, KNN, SVM, BPNN, ELM, MFO-ELM, GWO-ELM, and IMGWO+ELM have an accuracy of 92.31%, 92.69%, 93.54%, 93.85%, 93.08%,90.62% and 95.38% respectively. The comparison Figure 7-8 shows that at least 33 features in PCA obtain 93.38%, 95.00% in both G1020 and ORIGA datasets. We achieved optimal outcomes by employing a combination of dimensionality reduction methods known as PCA + LDA, that is, 93.87% and 95.38% in G1020 and ORIGA accordingly with 27 features described in Table 3.Afterward, an assessment of the employed scheme’s effectiveness implemented using a 10×5 fold stratified cross-validation (SCV) methodology. This scheme underwent manual hyperparameter tuning in each iteration to grasp high-level features. This primary classifier in the classification methodology was ELM, and the weights and biases were optimized using the enhanced GWO optimization technique. Furthermore, the model underwent testing using another meta-heuristic optimization technique called GWO. The study incorporated traditional classifiers like, KNN, SVM, BPNN, GWO-ELM, and IMGWO+ELM. A sample of 20 individuals was used, with a maximum iteration set at 100 for each classifier. The results obtained from a 10×5 fold cross-validation, incorporating ten interactions, are showcased in the Table 4 - 7. To observe the training sample’s performance convergence in a single execution for visual representation in Figure 9. The confusion matrix of the deployed model shown in Figure 10. The experimental evaluation showed that the results from our implemented approach exhibit superior classification performance compared to existing models while using a reduced number of features. The experimental findings suggest that the proposed MFO-ELM, GWO+ELM, and IMGWO+ELM models achieve lower condition and norm values, leading to improved accuracy compared to the traditional ELM approach. Table 8 and visulaized in Figures 11 and 12 compares classifier performance with our deployed work, named FCDT-WRP+PCA+LDA+IMGWO+ELM, is presented using various evaluation metrics like accuracy, sensitivity, and specificity. The findings suggest that our suggested model surpasses alternative classifiers in performance on both datasets. The Performance analysis of Proposed model with existing CAD models with G1020 and ORIGA datasets shown in Table 9.

**Table 2:**
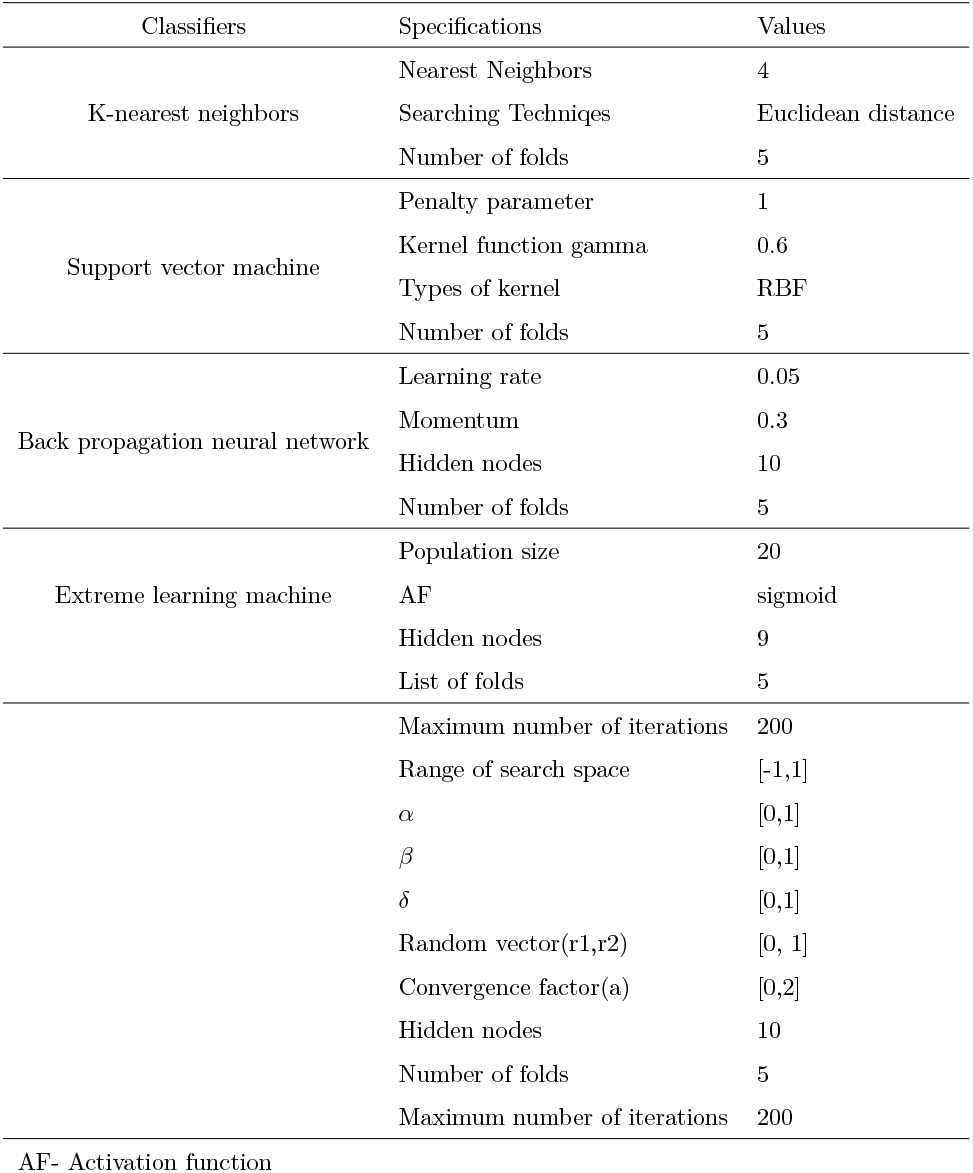
Compilation of hyperparameter specifications for distinct classifiers

**Table 3:** Comparative analyses (%) of the deployed model based on PCA with LDA techniques

| Proposed Method | No. of Features | G1020 |  |  | No. of Features | ORIGA |  |  |
| --- | --- | --- | --- | --- | --- | --- | --- | --- |
|  |  | Acc | Sen | Spe |  | Acc | Sen | Spe |
| PCA+GWO+ELM | 33 | 93.38 | 88.98 | 95.17 | 33 | 95.00 | 89.55 | 96.89 |
| **FDCT-WRP+PCA+LDA+MOD-GWO+ELM** | **27** | **93.87** | **89.83** | **95.52** | **27** | **95.38** | **91.04** | **96.89** |

Acc- Accuracy, Sen- Sensitivity, Spe-Specificity

**Table 4:**
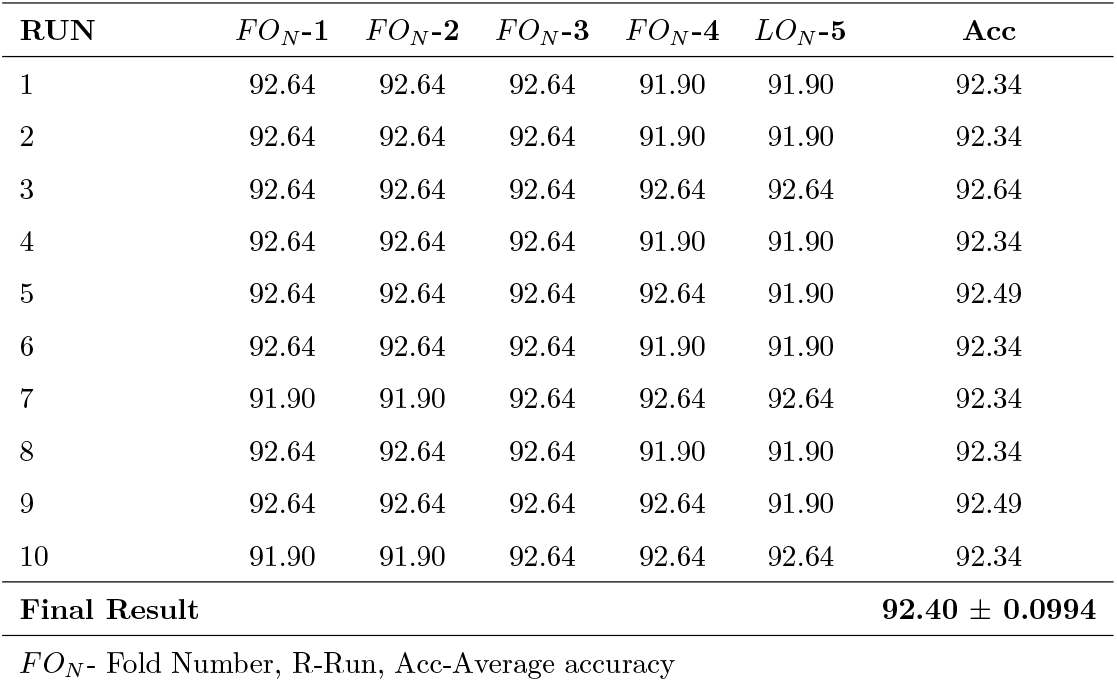
The effectiveness of the deployed CAD model using the Retinal G1020 dataset result(%) in 10×5-fold approaches with GWO+ELM

| **RUN** | *FO<sub>N</sub>-1* | *FO<sub>N</sub>-2* | *FO<sub>N</sub>-3* | *FO<sub>N</sub>-4* | *LO<sub>N</sub>-5* | **Acc** |
| --- | --- | --- | --- | --- | --- | --- |
| 1 | 92.64 | 92.64 | 92.64 | 91.90 | 91.90 | 92.34 |
| 2 | 92.64 | 92.64 | 92.64 | 91.90 | 91.90 | 92.34 |
| 3 | 92.64 | 92.64 | 92.64 | 92.64 | 92.64 | 92.64 |
| 4 | 92.64 | 92.64 | 92.64 | 91.90 | 91.90 | 92.34 |
| 5 | 92.64 | 92.64 | 92.64 | 92.64 | 91.90 | 92.49 |
| 6 | 92.64 | 92.64 | 92.64 | 91.90 | 91.90 | 92.34 |
| 7 | 91.90 | 91.90 | 92.64 | 92.64 | 92.64 | 92.34 |
| 8 | 92.64 | 92.64 | 92.64 | 91.90 | 91.90 | 92.34 |
| 9 | 92.64 | 92.64 | 92.64 | 92.64 | 91.90 | 92.49 |
| 10 | 91.90 | 91.90 | 92.64 | 92.64 | 92.64 | 92.34 |
| **Final Result** |  |  |  |  |  | **92.40 ± 0.0994** |

*FO<sub>N</sub>*- Fold Number, R-Run, Acc-Average accuracy

**Table 5:**
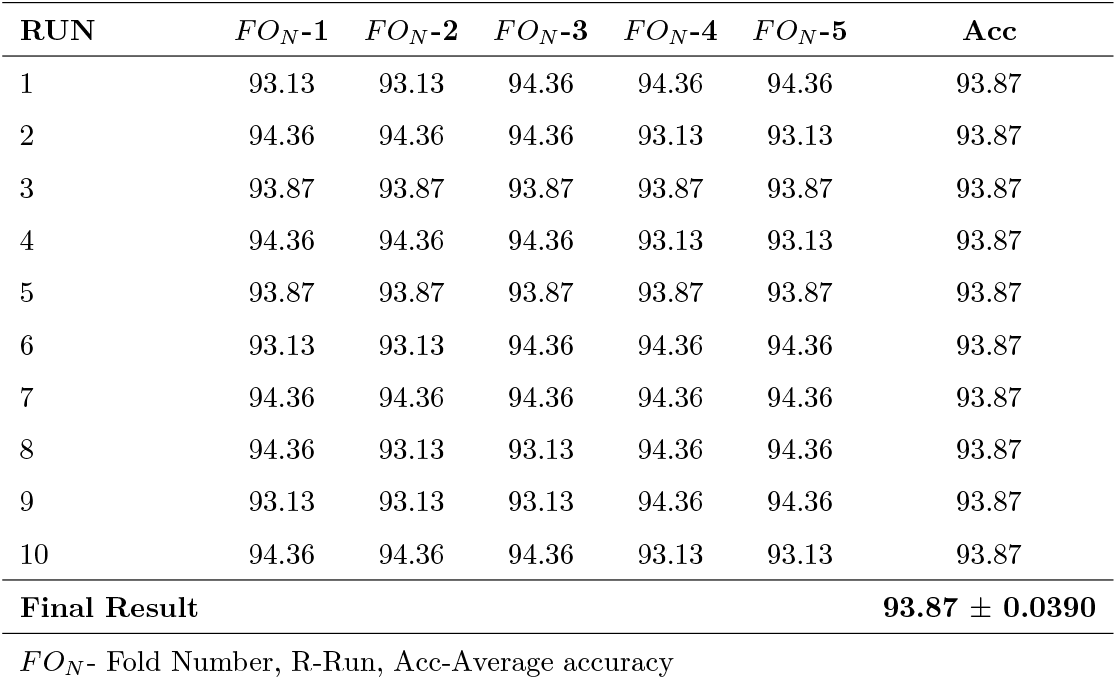
The effectiveness of the suggested CAD model using the Retinal G1020 datase result(%) in 10× 5-fold approches with IMGWO+ELM

**Table 6:** The effectiveness of the suggested CAD model using the Retinal ORIGA dataset result(%) in 10× 5-fold approches with GWO+ELM

| **RUN** | *FO<sub>N</sub>-1* | *FO<sub>N</sub>-2* | *FO<sub>N</sub>-3* | *FO<sub>N</sub>- 4* | *FO<sub>N</sub>-5* | **Acc** |
| --- | --- | --- | --- | --- | --- | --- |
| 1 | 94.81 | 94.81 | 93.86 | 93.86 | 93.86 | 94.24 |
| 2 | 94.81 | 94.81 | 93.86 | 93.86 | 93.86 | 94.24 |
| 3 | 94.81 | 94.81 | 94.81 | 94.81 | 94.81 | 94.81 |
| 4 | 94.81 | 94.81 | 94.81 | 93.86 | 93.86 | 94.43 |
| 5 | 94.81 | 94.81 | 94.81 | 93.86 | 93.86 | 93.43 |
| 6 | 94.81 | 94.81 | 94.81 | 94.81 | 94.81 | 94.81 |
| 7 | 94.81 | 94.81 | 94.81 | 94.81 | 94.81 | 94.81 |
| 8 | 94.81 | 94.81 | 94.81 | 94.81 | 94.81 | 94.81 |
| 9 | 94.81 | 94.81 | 94.81 | 94.81 | 94.81 | 94.81 |
| 10 | 94.81 | 94.81 | 94.81 | 94.81 | 94.81 | 94.81 |
| **Final Result** |  |  |  |  |  | **94.62 ± 0.2467** |
| *FO<sub>N</sub>*- Fold Number, R-Run, Acc-Average accuracy |  |  |  |  |  |  |

**Table 7:** The effectiveness of the suggested CAD model using the Retinal ORIGA dataset result(%) in 10× 5-fold approches with INGWO+ELM

| **RUN** | *FO<sub>N</sub>-1* | *FO<sub>N</sub>-2* | *FO<sub>N</sub>-3* | *FO<sub>N</sub>- 4* | *FO<sub>N</sub>-5* | **Acc** |
| --- | --- | --- | --- | --- | --- | --- |
| 1 | 95.76 | 95.76 | 95.76 | 95.10 | 95.10 | 95.50 |
| 2 | 95.00 | 95.00 | 95.76 | 95.76 | 95.76 | 95.46 |
| 3 | 95.10 | 95.10 | 95.10 | 95.10 | 95.10 | 95.10 |
| 4 | 95.00 | 95.00 | 95.76 | 95.76 | 95.76 | 95.46 |
| 5 | 95.00 | 95.00 | 95.76 | 95.76 | 95.76 | 96.46 |
| 6 | 95.10 | 95.10 | 95.10 | 95.10 | 95.10 | 95.46 |
| 7 | 95.00 | 95.00 | 95.76 | 95.76 | 95.76 | 95.46 |
| 8 | 95.00 | 95.00 | 95.76 | 95.76 | 95.76 | 95.46 |
| 9 | 95.10 | 95.10 | 95.10 | 95.10 | 95.10 | 95.10 |
| 10 | 95.76 | 95.76 | 95.76 | 95.76 | 95.76 | 95.76 |
| **Final Result** |  |  |  |  |  | **95.38 ± 0.2063** |
| *FO<sub>N</sub>*- Fold Number, R-Run, Acc-Average accuracy |  |  |  |  |  |  |

**Table 8:** Comparative Analysis (%) employed scheme Glaucoma datasets with different specifications

| Optimization Method+Classifiers | G1020 Dataset |  |  | ORIGA Dataset |  |  |
| --- | --- | --- | --- | --- | --- | --- |
|  | Acc | Sen | Spe | Acc | Sen | Spe |
| KNN | 90.69 | 84.75 | 93.10 | 92.31 | 85.07 | 94.82 |
| SVM | 91.67 | 86.44 | 93.79 | 93.08 | 88.06 | 94.82 |
| BPNN | 92.65 | 87.29 | 94.83 | 93.85 | 92.54 | 94.30 |
| ELM | 93.63 | 88.98 | 95.52 | 93.46 | 88.06 | 95.34 |
| MFO+ ELM | 93.14 | 88.14 | 95.17 | 93.08 | 88.06 | 94.82 |
| GWO + ELM | 92.40 | 88.14 | 94.14 | 94.62 | 94.03 | 94.82 |
| **FDCT-WRP+PCA+LDA+IMGWO+ELM (Proposed Model)** | **93.87** | **89.83** | **95.52** | **95.38** | **91.04** | **96.89** |

Acc: Accuracy, Sen: Sensitivity , Spe: Specificity

**Table 9:** Performance analysis of Proposed model with existing CAD models with G1020 and ORIGA datasets

| Existing Methods | Acc (%) |  |
| --- | --- | --- |
|  | G1020 | ORIGA |
| TIA-Net (SOD+Attention) [79] | — | 85.70 |
| 2D-FBSE-EWT [80] | — | 91.01 |
| SMOTE + RF [81] | — | 78.30 |
| SMOTE + RF [81] | — | 82.80 |
| HOG +SVM [51] | 83.32 | — |
| HOG + PNN [51] | 87.92 | — |
| HOG + RNN [51] | 85.72 | — |
| **FDCT-WRP+PCA+LDA+IMGWO+ELM (Proposed Model)** | **93.87** | **95.38** |

Acc- Accuracy

**Figure 7.**
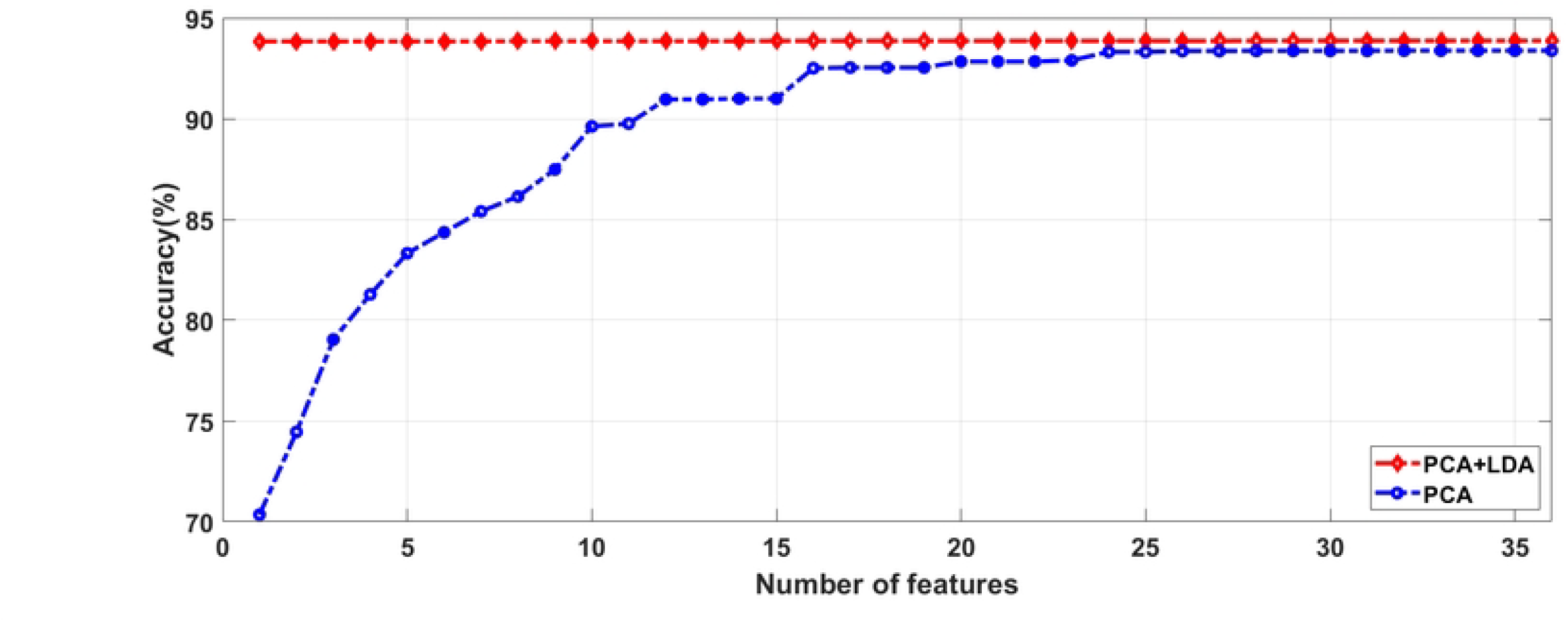
Accuracy with respect to number of Features using G1020 Dataset.

**Figure 8.**
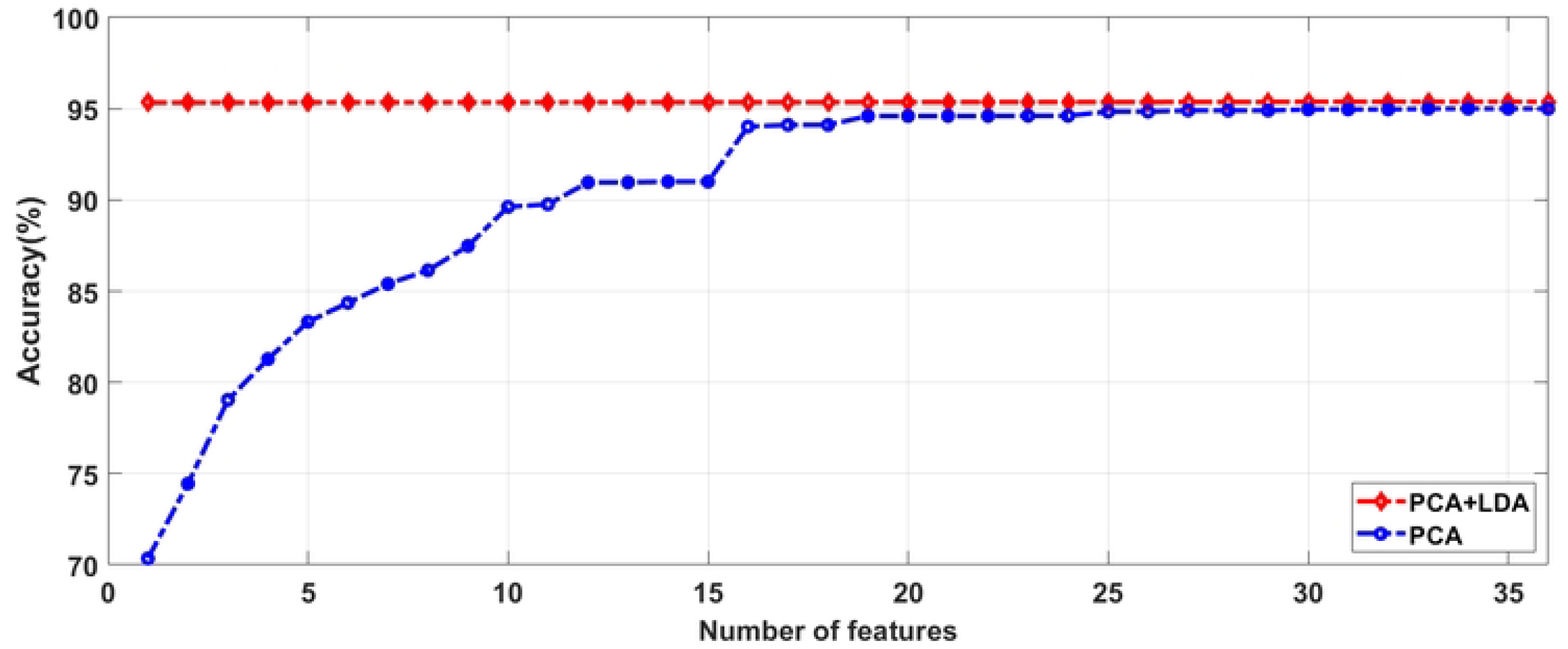
Accuracy with respect to Number of Features using ORIGA Dataset.

**Figure 9.**
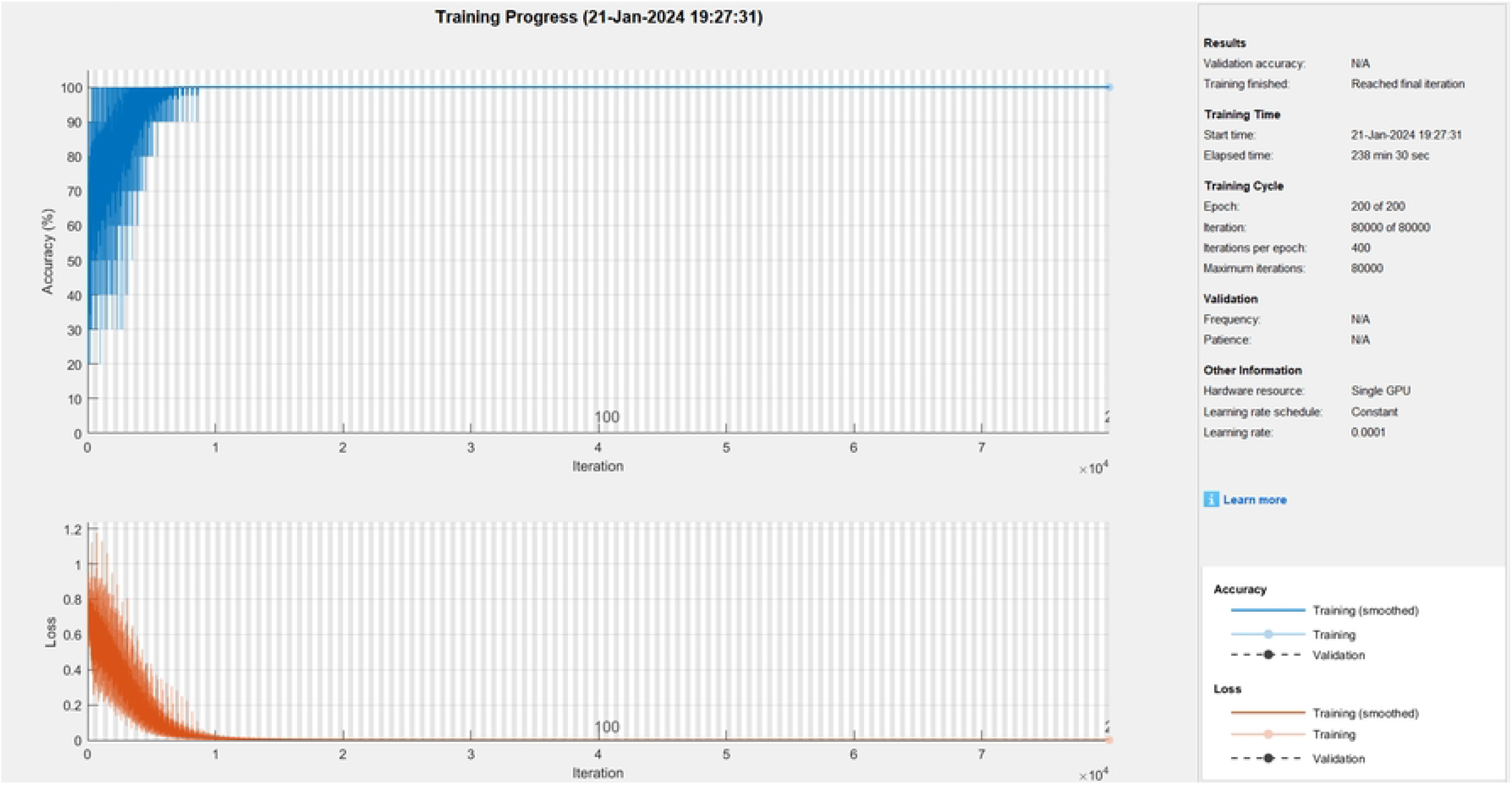
Accuracy with respect to Loss on one run using G1020 and ORIGA datasets

**Figure 10.**
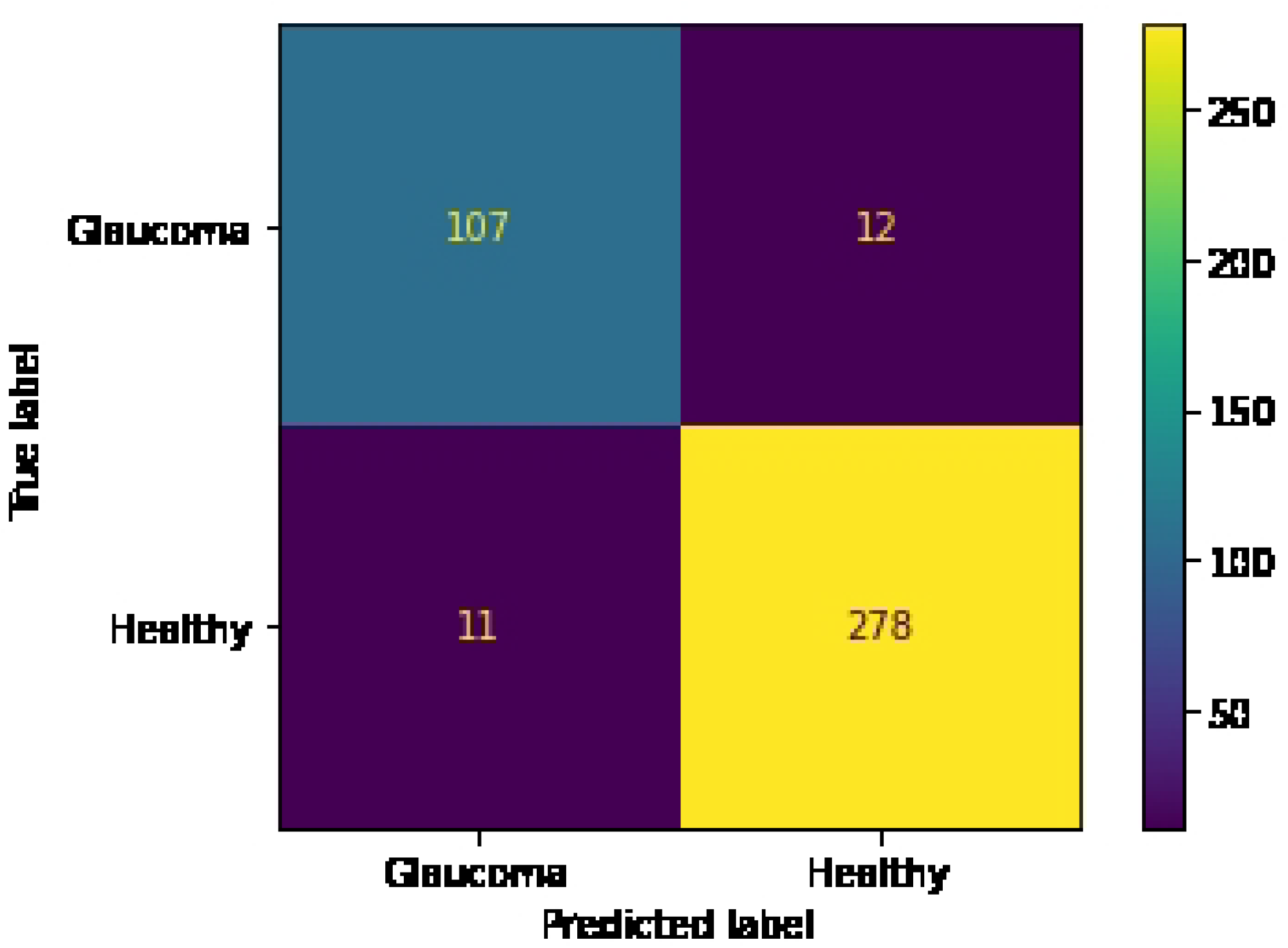
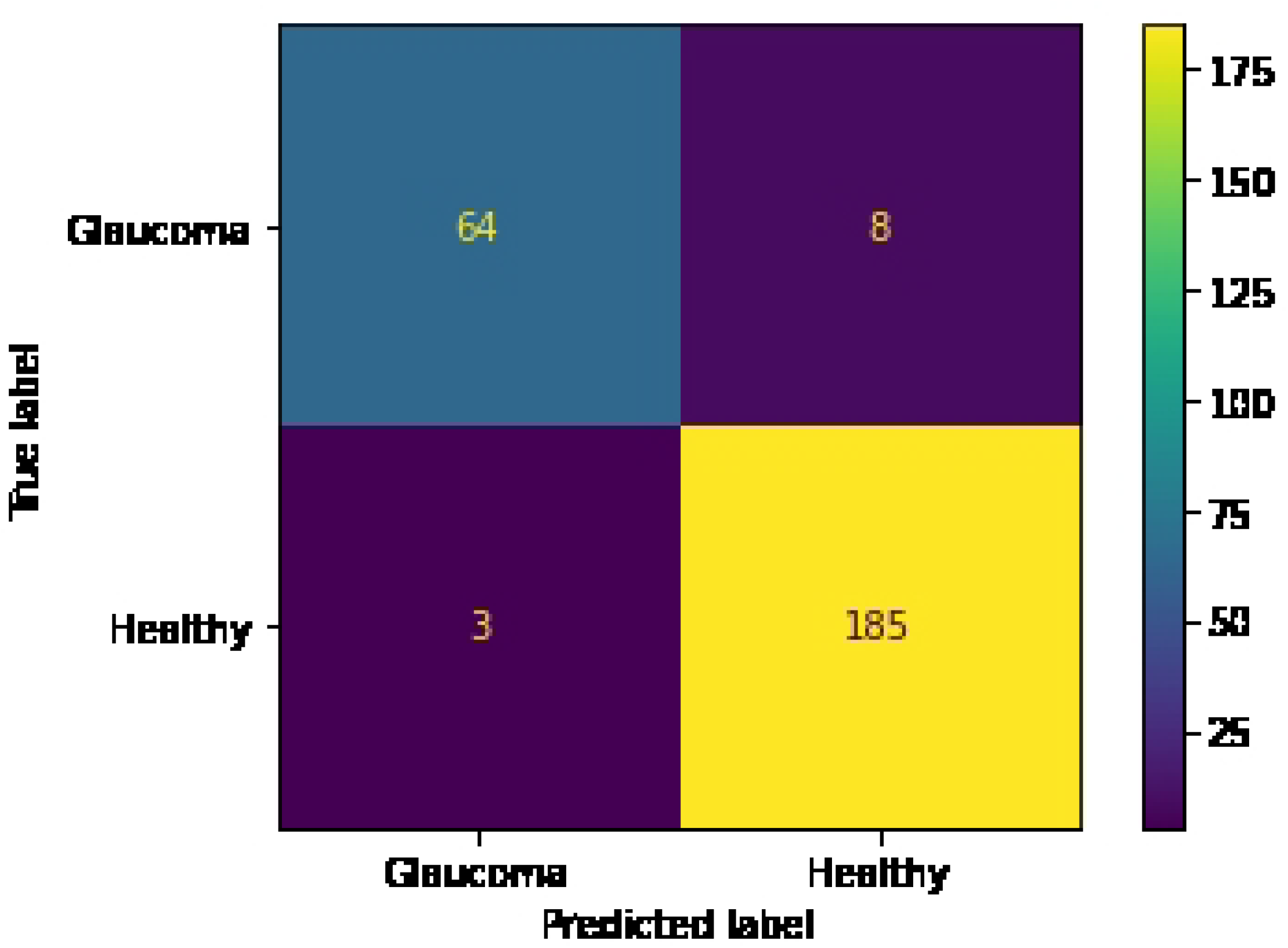
Confusion matrix of deployed scheme based on (a) G1020 and (b) ORIGA Datasets.

**Figure 11.**
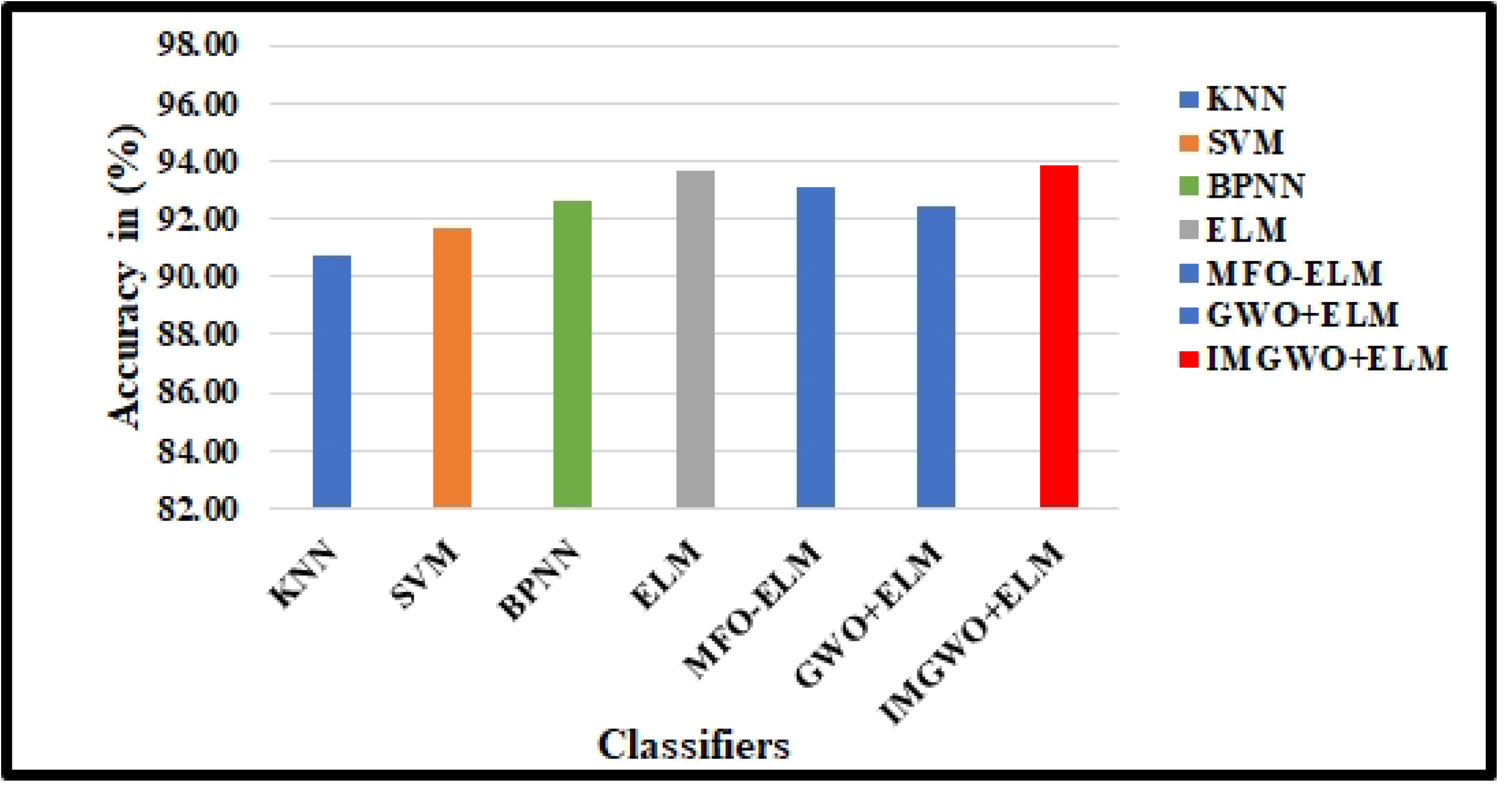
Performance Analysis on several classifiers with existing model using G1020 Dataset.

**Figure 12.**
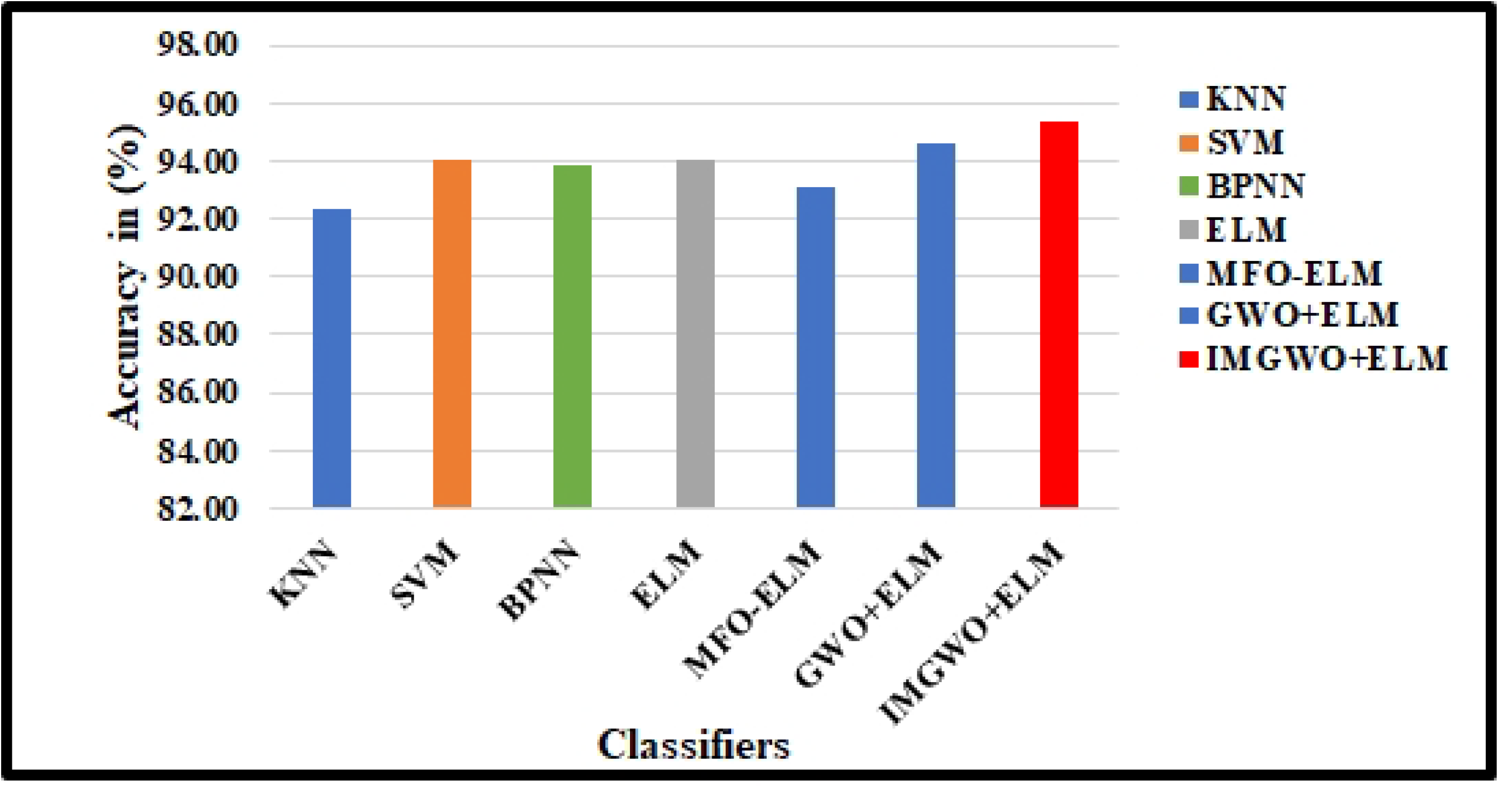
Performance Analysis on several classifiers with existing model using ORIGA Dataset.

#### 4.1. Advantages and disadvantages

Machine learning has widely used in various medical applications, such as biomedical image processing and analysis. Through the utilization of AI-boosted computer-aided design (CAD) models, ophthalmologists’ expertise has advanced to the extent that they can now detect and diagnose glaucoma more effectively, leading to improved outcomes and shorter diagnostic intervals. In applications increasingly crucial to healthcare, implement these models has been limited due to the lack of explainability. GlaucoXAI utilizes seven distinct gradient-based explanation techniques—VG, GBP, IG, GIG, Smooth-Grad, GCAM, and GGCAM—to enhance the transparency of ML approches. Each method of explainable artificial intelligence (XAI) possesses its own distinct characteristics and can offer assistance in various situations, each with its own set of advantages and drawbacks. For instance, VG is straightforward and benefits from compatibility with traditional machine learning platforms like TensorFlow [39] and PyTorch [45]. This implies that Vanilla Gradient (VG) can be used with any deep neural network without needing to change its architecture. However, the generated saliency maps by VG tend to be noisy, and they also lose the influence of features over time due to gradient saturation, as noted in previous studies. On the contrary, Gradient Backpropagation (GBP) is efficient to implement but has limitations; it only works with ML models for classification and does not produce visualization maps that distinctly represent different classes. In recent times, IG has gained popularity due to its simplicity in implementation, lack of network instrumentation requirement, and a fixed number of gradient calls. GIG is an improvement designed to address IG’s false perturbations issue, However, it necessitates making a decision at every stage from the starting point to the input, making the path direction variable. SmoothGrad, while aiding in enhancing visualizations of the genuine signal, lacks class discrimination ability, which is a significant limitation. In contrast, GCAM enables the interpretation of any ELM by emphasizing the distinctive region, aiding in the comprehension of its internal operations. GGCAM, a combination of GBP and GCAM benefits, was introduced to address the issue of lower-resolution heatmaps associated with GCAM. These methods of explanation underwent extensive examination in the context of two common brain imaging analysis tasks: assessing the severity of gliomas and pinpointing their locations. In both situations, detailed gradient-based maps like VG, GBP, IG, GIG, and SmoothGrad were employed to emphasize all pertinent characteristics, regardless of the specific class chosen, as depicted in Figure 2. Alternatively, GCAM and GGCAM identify the crucial areas guiding the network’s decision-making process. This aligns with research in [27], indicating that humans comprehend regions rather than individual pixels more effectively.

Our suggested CAD scheme utilized a fast discrete curvelet transform with wrapping technique for extracting curve-like characteristics from fundus images. This method captures 2D singularities, such as curves in retinal fundus images. An integrated feature reduction technique, the PCA + LDA approach, simplifies identifying pertinent features. The advantages of the IMGWO+ELM method include a concise network architecture, heightened condition value, and superior generalization outcomes achieved through accelerated learning.

Finally, there are several shortcomings associated with the suggested method, as detailed below. The validation of the proposed computer-aided design (CAD) model has been restricted to retinal glaucoma fundus images. Although the model’s application to dual classifications has been scrutinized, forthcoming research will concentrate on broadening its applicability to address challenges in multi-class classification. Likewise, the proposed IMGWO algorithm necessitates adjustments to multiple parameters for optimization. Subsequent versions might utilize an enhanced optimization method that involves a reduced number of features.

## 5. Conclusion and Future work

Our proposed manuscript presented an improved expalinability framework, called Glaucoma explainable artificial intelligence) GlaucoXAI provides support in understanding the behavior of deep learning networks by employing cutting-edge visualization techniques such as attention maps. As a post hoc tool, GlaucoXAI can be applied to any already existing deep neural models, providing significant insights into how they operate. Our two case studies underscore the importance of integrating XAI techniques in medical image analysis. Furthermore, GlaucoXAI facilitates ELM classifier for glaucoma detection. Our findings emphasize the pivotal role of XAI in medical imaging tasks, aiding in the comprehension of ML models and expediting their adoption by medical professionals. This paper introduces a sophisticated computer-aided design (CAD) model designed for the categorization of glaucoma or healthy images. The model adeptly identifies pertinent features in fundus images by utilizing a fast discrete curvelet transform with a wrapping (FDCT-WRP) process. A combined PCA + LDA technique is applied to further enhance feature reduction, resulting in reduced and more prominent features. Subsequently, the CAD model employs the IMGWO-ELM, a faster learning algorithm, to train the SLFN. The CAD model’s classification performance is rigorously evaluated across two standard fundus image datasets. The experimental results reveal that the proposed CAD model achieves superior classification performance with fewer features than existing models. In future research, the efficacy of employed will be tested for generalization across various imaging modalities. Another potential avenue is the exploration of hybridizing ELM with a less parameter-based optimization algorithm, assessing its effectiveness in solving multi-class classification tasks. Additionally, the paper suggests considering deep learning algorithms as potential alternatives to the proposed model. In future research, our attention will be directed towards quantitatively evaluating XAI methods. This evaluation aims to gauge the effectiveness of sensitivity maps generated by these methods and explore their correlation with deep learning accuracy metrics. We plan to conduct further experiments in the realm of multi-modal glaucoma-guided opthamalogists to enhance our understanding. Additionally, we aim to delve into the potential of extracting quantitative features, such as tumor volume and centroid, from explanation methods.

## Ethical declarations

*Ethics approval and consent to participate*

Not applicable.

*Clinical trial number*

Not applicable.

*Institutional Review Board Statement*

Not applicable.

*Informed Consent Statement*

Not applicable.

## Data Availability Statement

The datasets analysed during the current study are available in G1020 [77] and ORIGA [78].

## Authors Contribution

D.M., S.K.S., S.D. worked on Conceptualization, Data curation, Experiment, Methodology, Resources, Validation, Writing – original draft. B.L., S.M. performed Investigation, Project administration, Supervision, Writing – review & editing.

